# A tonic PDGFA-dependent niche organizer maintains alveolar homeostasis

**DOI:** 10.64898/2026.10.02.756373

**Authors:** Rebecca Shelley Frabotta, Arvind Konkimalla, Thomas Whitlow, Kyoshiro Tsuge, Shuyu Liu, Michael T. Forrester, Aleksandra Tata, Purushothama Rao Tata

**Affiliations:** Department of Cell Biology, Duke University School of Medicine, Durham, NC 27710, USA; Duke University Cell and Molecular Biology (CMB) Graduate Program; Division of Pulmonary, Allergy and Critical Care Medicine, Duke University School of Medicine, Durham NC 27710, USA; Department of Surgery, Surgical Sciences, Duke University School of Medicine, Durham, NC 27710, USA; Duke Regeneration Center, Duke University, Durham, NC 27710, USA; Duke Cancer Institute, Duke University School of Medicine, Durham, Durham, NC 27710, USA; Center for Advanced Genomic Technologies, Duke University, Durham, NC 27710, USA

## Abstract

Tissue homeostasis requires mechanisms that preserve quiescence while allowing rapid activation after injury, but how tissues actively restrain injury responses during homeostasis remains poorly understood. Here we identify alveolar type 1 (AT1) cells as organizers of the distal lung niche through tonic expression of PDGFA, which maintains alveolar fibroblast identity and restrains myofibroblast activation. Cell type-specific ablation revealed that loss of AT1s, but not other major alveolar cell populations, was sufficient to evoke spontaneous fibroblast activation and tissue-wide responses that recapitulate injury. In silico ligand-receptor analysis, ex vivo screening, and conditional genetic models identified AT1-derived PDGFA as a key homeostatic signal. Loss of AT1-derived PDGFA caused spontaneous myofibroblast induction and tissue remodeling, whereas increasing PDGFA promoted fibroblast proliferation while suppressing myofibroblast differentiation. Integrated phosphoproteomic, transcriptomic, and chromatin accessibility analyses showed that PDGFA engages SHP2-ERK signaling, suppresses MRTFA-associated SRF programs, and promotes E2F-associated cell-cycle programs, thereby opposing the pro-fibrotic effects of TGFβ-driven fibroblast activation. Restoring PDGFA signaling abrogates myofibroblast features from human idiopathic pulmonary fibrotic lungs and attenuated fibrotic remodeling in vivo. These findings define a tonic AT1-derived PDGFA signal that maintains alveolar homeostasis and offers a therapeutic target to prevent or revert lung fibrosis.

## Introduction

Maintaining tissue homeostasis despite environmental and physiological perturbations is a fundamental property of living systems ^1–4^. The strategies that achieve homeostasis vary across organs: rapidly renewing tissues such as the intestine and epidermis rely on continuous progenitor activity, whereas tissues such as the adult lung remain relatively quiescent at steady state but can rapidly activate regenerative programs in response to injury ^2,5–8^. Despite extensive work defining pathways that activate regenerative programs, how tissues actively maintain homeostasis, whether through cellular quiescence or continuous renewal, remains poorly understood ^2,8,9^. A key unresolved question is whether tissue homeostasis emerges from distributed interactions among many cell types or is actively coordinated by specific cell populations that function as tissue organizers ^10–14^. Such organizers could integrate local environmental cues and transmit instructive signals to neighboring cells, thereby setting thresholds for activation, calibrating repair, and promoting return to quiescence after injury. Molecules that enforce this control are uniquely positioned to function as homeostatic checkpoints, restraining inappropriate activation under baseline while enabling rapid entry into regenerative states following tissue injury ^15,16^.

In the distal lung, the epithelium, endothelium, fibroblasts, and other resident cells of the alveoli exhibit remarkable quiescence, yet all have been shown to respond to injury and participate in the repair process ^6,7,17–19^. Although the precise cellular response varies with the nature and severity of injury, repair engages a broadly conserved cascade of cellular state transitions ^20–24^. For example, otherwise quiescent fibroblasts rapidly respond to injury by transitioning through primed and inflammatory states before adopting myofibroblast and remodeling programs, while activated alveolar type-2 (AT2) cells pass through transitional states en route to specialized alveolar type-1 (AT1) differentiation ^20,21,24,25^. However, how these injury-responsive cell populations are actively maintained in a quiescent state at baseline, and whether a specific alveolar cell type serves as a homeostatic organizer of this multicellular niche, remains unclear.

Here, using cell ablation, niche communication assays, conditional genetic models, and integrated phosphoproteomic, transcriptomic, and chromatin accessibility analyses, we identify PDGFA derived from AT1s as a critical signal that acts on alveolar interstitial fibroblasts to maintain their quiescent state. Loss of AT1s or AT1-derived PDGFA alone was sufficient to trigger an injury-like response in otherwise uninjured alveolar tissue, characterized by rapid fibroblast activation, myofibroblast differentiation, and an organ-wide fibrotic response. Together, our findings identify AT1s as a tissue organizer that couples environmental sensing to epithelial-stromal communication, thereby maintaining fibroblast quiescence and coordinating the transition from homeostasis to repair and its subsequent resolution.

## Results

### Cell ablation models reveal AT1 cells are alveolar niche organizers and maintains tissue homeostasis

To identify alveolar cell populations that function as homeostatic organizers, we performed cell type-specific ablation of major epithelial, stromal, immune, and endothelial populations and assessed the resulting tissue responses. Previous studies have used AT1 ablation models and revealed that loss of AT1s alone is sufficient to invoke injury responses in the lung ^21^. However, whether AT1s are uniquely able, as opposed to loss of any other cell type, to induce such injury-repair response has not been established. To test this, we assembled a panel of cell type-specific ablation mouse models targeting AT1s (*Ager-creER;R26R-DTR*), AT2s (*Sftpc-creER;R26R-DTA-tdT*), fibroblasts (*Pdgfra-creER;R26R-DTR*), macrophages (*Cd169-DTR*), and endothelial cells (*Cdh5-creER;R26R-DTR*) (Figure 1a). We verified ablation efficiency in each model by quantifying the corresponding lineage markers, including TCF21, SFTPC, F4/80 to identify fibroblasts, AT2s, and macrophages, respectively (Extended Data Figure 1a–c). At 5-7 days after ablation, we assessed alveolar fibroblast cell states by immunostaining for PDGFRA, ACTA2, and LEL (*Lycopersicon esculentum* lectin, a marker for AT1s and alveolar architecture) (Figure 1b). Strikingly, among the cell populations tested, only AT1 cell ablation elicited robust activation of alveolar fibroblasts, marked by widespread induction of ACTA2 (Figure 1b and Extended Data Figure 1d). Quantification of ACTA2^+^ area confirmed a significant increase following AT1 cell ablation compared with untreated wild-type controls, whereas ablation of AT2s, fibroblasts, macrophages, or endothelial cells produced no significant increase in ACTA2^+^ myofibroblasts (Figure 1c). These findings demonstrate that loss of AT1s, but not the other major alveolar cell populations tested, is sufficient to trigger spontaneous fibroblast activation, suggesting that AT1s serve as a niche organizer and maintains the quiescent state of the alveolar niche.

**Figure 1:**
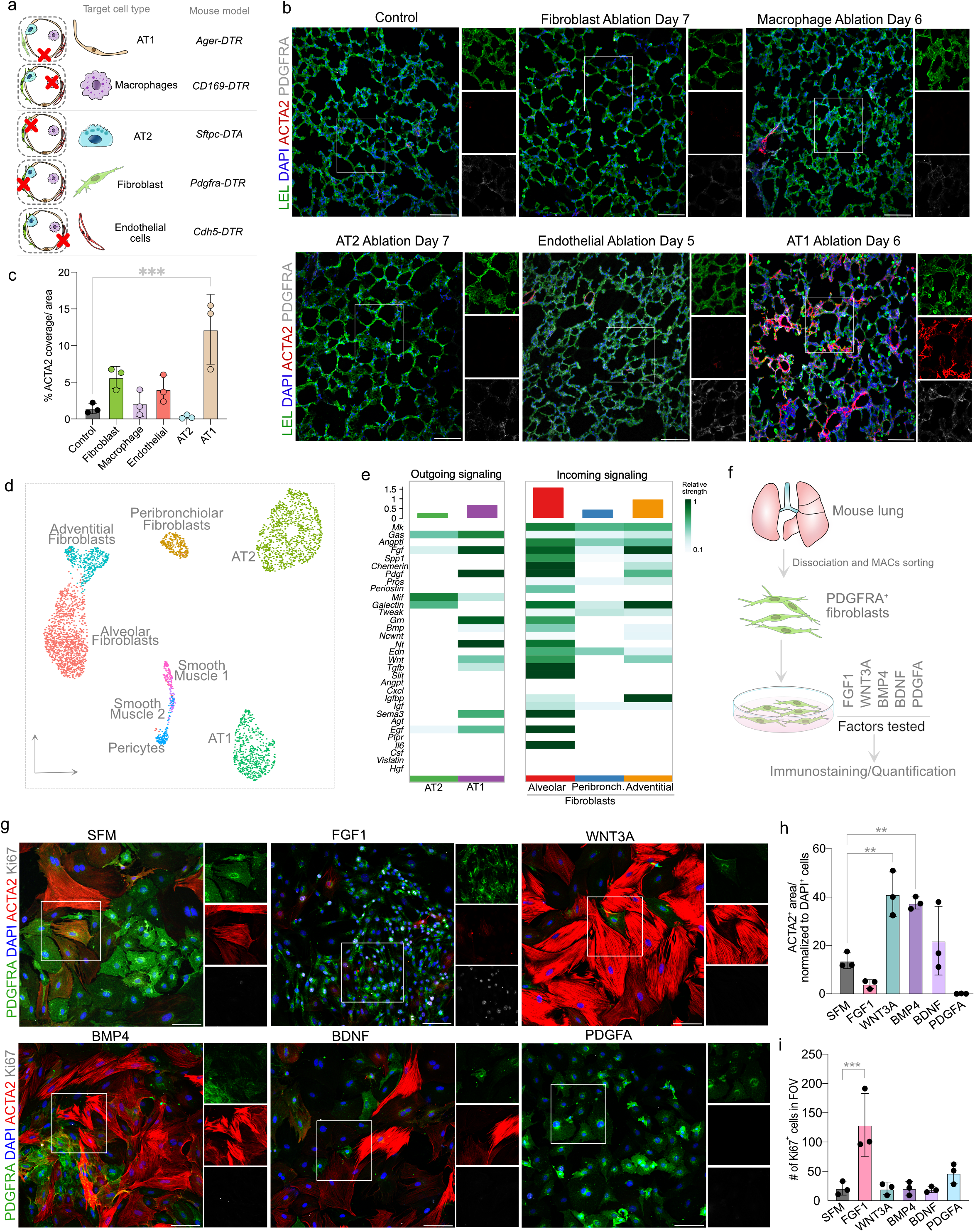
AT1 cell loss initiates alveolar fibroblast activation through PDGFA signaling. a. Schematic of mouse models used for ablation of indicated cell types in the alveoli. b. Immunostaining for LEL (green), ACTA2 (red), and PDGFRA (grey) in control and specific cell type ablated lungs. Scale bar: 100μm. c. Quantification of % ACTA2 coverage/area in control and ablation models. n=3 mice per condition. One-way ANOVA with Dunnett’s multiple comparisons test vs. control; ***p = 0.0004. d. UMAP visualization of mouse lung mesenchymal and epithelial cell populations from scRNA-seq data. e. Heatmaps of relative signaling strength for outgoing (left) and incoming (right) signaling pathways across cell types, calculated using CellChat. Bar plots above each heatmap indicate overall relative signaling strength per cell type; heatmap color intensity denotes relative pathway strength for each cell type-pathway pair. f. Schematic of AT1-derived molecules tested in alveolar fibroblast cultures. g. Immunostaining for PDGFRA (green), ACTA2 (red), and Ki67 (grey) in primary alveolar fibroblast cultures collected at day-3 after treatment. Serum-free media (SFM) control verses FGF1 (10ng/ml) + Heparin (10ug/ml), WNT3A (5ng/ml), BMP4 (50ng/ml), BDNF (50ng/ml), and PDGFA (10ng/ml). Scale bar: 100μm. h. Quantification of ACTA2^+^ area/normalized to DAPI count in FOV. One-way ANOVA with Dunnett’s multiple comparisons test vs. SFM; **p<0.01. n=3 mice per condition. i. Quantification of the number of Ki67^+^ cells in FOV. One-way ANOVA with Dunnett’s multiple comparisons test vs. SFM; ***p = 0.0005. n=3 mice per condition. DAPI stains nuclei in all panels. Statistical significance is indicated by asterics. In all graphs data are presented as mean ± SD.

### In silico and ex vivo screens identify AT1-derived PDGFA as a regulator of fibroblast identity

To identify molecular signals through which AT1s regulate alveolar fibroblasts, we applied CellChat, a computational framework for inferring ligand-receptor-mediated cell-cell communication from single-cell RNA sequencing (scRNA-seq) data ^26^. Analysis of control mouse lungs predicted extensive communication among alveolar epithelial cells, fibroblasts, pericytes, and smooth muscle cells. Among the predicted AT1-to-alveolar fibroblast interactions were *Pdgfa-Pdgfra*, *Fgf1-Fgfr1/2/3/4*, *Bdnf-Ntrk*, *Bmp4-Bmpr1a/Acvr2a*, *Wnt3a-Fzd/Lrp*, and *Hbegf-Egfr* signaling (Figure 1d-e). Visualization of ligand and receptor expression confirmed their respective enrichment in AT1s and alveolar fibroblasts, while quantitative analysis of signaling patterns revealed prominent outgoing activity from AT1s and incoming activity in alveolar fibroblasts (Extended Data Figure 2a,b). CellChat analysis of human lung scRNA-seq data similarly predicted communication between these cell populations and identified signaling pathways conserved across mouse and human lungs (Figure 1d-e and Extended Data Figure 2c-f). Based on cross-species conservation and enrichment of ligand expression in AT1s, we prioritized FGF1, PDGFA, BMP4, WNT3A, and BDNF for functional testing (Figure 1f).

We next tested the effects of candidate AT1-derived trophic factors on alveolar fibroblasts. Because standard fibroblast culture conditions promote spontaneous activation and myofibroblast differentiation, we established a chemically defined, serum-free medium (SFM) in which fibroblasts exhibited a low, measurable level of spontaneous myofibroblast differentiation, as assessed by ACTA2 expression. This differentiation was markedly enhanced by supplementation with 10% serum (Figure 1f,g and Extended Data Figure 2g). We therefore used SFM as a baseline condition to quantitatively assess the effects of AT1-derived trophic factors on fibroblast differentiation and proliferation. Purified primary alveolar fibroblasts were cultured for 3-days in SFM supplemented with FGF1, WNT3A, BMP4, BDNF, or PDGFA, and myofibroblast differentiation and proliferation were assessed by immunostaining for ACTA2 and Ki67, respectively (Figure 1f,g).

This screen revealed three distinct responses. First, FGF1 and PDGFA most strongly suppressed ACTA2 expression relative to SFM alone (Figure 1g-h). However, FGF1 significantly increased the number of Ki67^+^ fibroblasts while suppressing ACTA2 expression (Figure 1g-i). Second, WNT3A and BMP4 unexpectedly increased ACTA2 expression relative to SFM controls, reaching levels comparable to those observed in SFM supplemented with 10% serum (Figure 1g-h and Extended Data Figure 2g). In contrast, PDGFA strongly suppressed ACTA2 expression without inducing overt fibroblast proliferation (Figure 1g-i). Quantification of ACTA2 expression and Ki67^+^ cells therefore identified PDGFA as the only candidate tested that robustly restrained myofibroblast differentiation without concomitantly promoting proliferation. Together, these findings identify AT1-derived PDGFA as a candidate homeostatic signal that preserves alveolar fibroblast identity and suppresses acquisition of an activated myofibroblast state.

### AT1-derived PDGFA is required for fibroblast homeostasis in vivo

Having identified PDGFA as a candidate AT1-derived trophic factor that regulates fibroblast state ex vivo, we next asked whether AT1-derived PDGFA is required to maintain fibroblast quiescence in vivo. We first performed targeted CellChat analysis of PDGFA-PDGFRA signaling in mouse and human lung scRNA-seq datasets. Although AT1s showed predicted PDGFA-mediated communication with multiple fibroblast populations, the strongest interaction was observed with alveolar fibroblasts (Figure 2a and Extended Data Figure 3a,b). To validate AT1s as a source of PDGFA in vivo, we performed proximity ligation in situ hybridization (PLISH) on lung sections. Co-detection of *Pdgfa* and the AT1 marker *Ager* demonstrated enrichment of *Pdgfa* transcripts in AT1s (Figure 2b). The specificity of the PLISH probes used to detect *Pdgfa* was further validated using AT1s derived from *Pdgfa^fl/fl^* mice (Extended Data Figure 3c). Analysis of scRNA-seq data spanning AT2s and their derivatives, including proliferating AT2s, early and late transitional states, and mature AT1s, further revealed a progressive increase in *Pdgfa* expression beginning in transitional states and reaching its highest levels in AT1s (Figure 2c).

**Figure 2:**
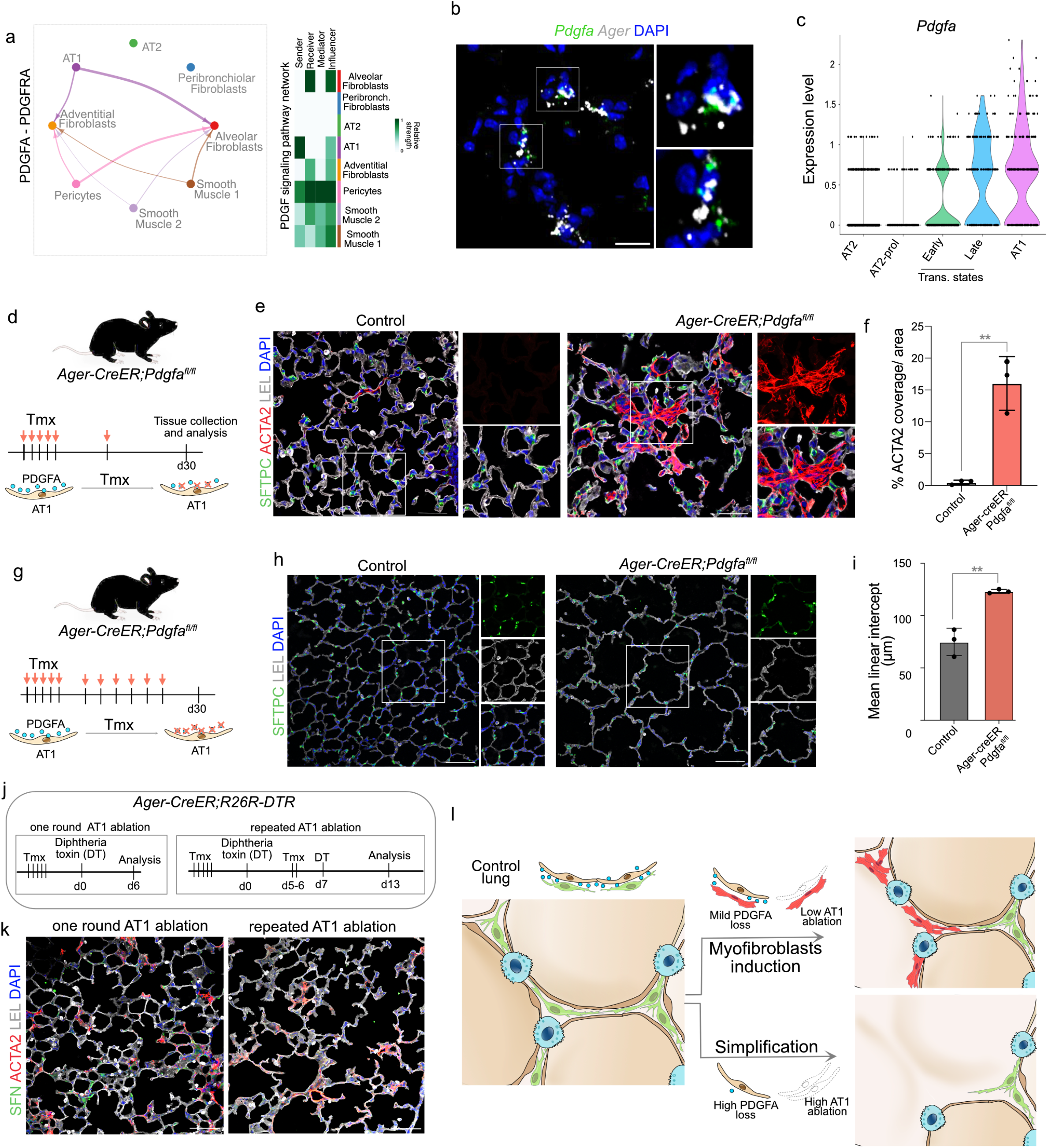
AT1-derived PDGFA maintains alveolar fibroblasts identity *in vivo*. a. Chord diagram depicting predicted PDGFA-PDGFRA ligand-receptor signaling between cell types (top), with arrow color and width indicating signaling source and relative strength. Heatmap (right panel) shows the relative importance of each cell type in the PDGF signaling pathway network across four roles: sender, receiver, mediator, and influencer. b. Proximity ligation in situ hybridization (PLISH) for *Pdgfa* (green) and *Ager* (grey) in control mouse alveolar tissue. Scale bar: 20μm. c. *Pdgfa* expression across epithelial cell populations in mouse lung scRNA-seq data shown as log-normalized expression. Cell populations include AT2, proliferating AT2 (AT2-prol), early and late epithelial transitional states, and AT1 cells. d. Schematic depicting loss of AT1-derived PDGFA using *Ager-creER;Pdgfa^fl/fl^* mouse model. Sequential administration of tamoxifen (TMX) for achieve low Pdgfa deletion. e. Immunostaining for SFTPC (green), ACTA2 (red), and LEL (grey) in the short TMX timeline administration model. Scale bar: 50μm. f. Quantification of ACTA2 coverage. Unpaired two-tailed t test; **p = 0.0031. n=3 mice per condition. g. Schematic depicting loss of AT1-derived PDGFA using *Ager-creER;Pdgfa^fl/fl^* mouse model. Sequential administration of tamoxifen (TMX) for achieve high Pdgfa deletion. h. Immunostaining for SFTPC (green) and LEL (grey) in the long TMX timeline administration model. Scale bar: 100μm. i. Quantification of MLI in the long TMX timeline. Unpaired two-tailed t test; **p=0.0032. n=3 mice per condition. j. Schematic depicting AT1 cell ablation using *Ager-creER;R26R-DTR* mouse model. Sequential administration of tamoxifen (TMX) and diphtheria toxin (DT) followed by tissue collection immunofluorescence analysis. k. Immunostaining for SFN (green), ACTA2 (red), and LEL (grey) in both single and repeated AT1 ablation models. Scale bar: 100μm. l. Schematic representation for experimental outcomes illustrating emergence of myofibroblasts in AT1 ablation and low deletion Pdgfa deletion model and tissue simplification in repeated AT1 ablation and high Pdgfa deletion model. DAPI stains nuclei in all panels. Statistical significance is indicated by asterics. In all graphs data are presented as mean ± SD.

To determine whether AT1-derived PDGFA is required to maintain fibroblast quiescence in vivo, we generated *Ager-CreER;Pdgfa^fl/fl^* mice and used low frequency tamoxifen regimens to achieve lower levels of *Pdgfa* deletion in AT1s, followed by tissue analysis 30 days later (Figure 2d). ACTA2 immunostaining revealed a dose-dependent tissue response. Low-dose tamoxifen induced spontaneous accumulation of ACTA2^+^ cells in *Ager-creER;Pdgfa^fl/fl^*lungs but not in controls (Figure 2e), with quantification confirming a significant increase in ACTA2^+^ area (Figure 2f). These spontaneously emerging myofibroblasts also expressed additional activation-associated markers, including TIMP1 and CTHRC1, and were accompanied by increased LEL^+^ macrophages and extensive tissue remodeling resembling that observed following lung injury (Extended Data Figure 3d-f). Thus, partial loss of AT1-derived PDGFA was sufficient to elicit an injury-like response within the alveolar niche.

In contrast, high-dose tamoxifen treatment of *Ager-creER;Pdgfa^fl/fl^*mice (Figure 2g) resulted in marked alveolar simplification resembling an emphysema-like phenotype, which was confirmed by mean linear intercept analysis (Figure 2h-i). The distinct phenotypes produced by low and high levels of *Pdgfa* deletion suggested that the magnitude of AT1-derived PDGFA signaling may differentially regulate fibroblast properties and alveolar tissue organization. We therefore asked whether altering AT1 cell abundance itself would produce a similar graded response. To do so, we employed *Ager-CreER;R26R,DTR* mice and performed single or repeated AT1 ablation (Figure 2j). A single round of AT1 cell ablation resulted in abundant ACTA2^+^ cells within the alveolar interstitium, whereas two rounds of AT1 ablation led to pronounced alveolar simplification (Figure 2k). Together, these findings suggest that AT1 cell abundance or the levels of AT1-derived PDGFA ligands regulate alveolar fibroblast state and, consequently, alveolar tissue organization (Figure 2l).

### PDGFA maintains fibroblast identity in a dose-dependent manner and modulates TGFβ-driven activation

The preceding data indicate that AT1-derived PDGFA suppresses the differentiation of alveolar fibroblasts into myofibroblasts. This finding contrasts with the prevailing view that PDGFA promotes fibrosis ^27,28^. Indeed, PDGFA has been used as a component of pro-fibrotic cocktails together with TGFβ, TNFα, and LPA (^29^. To determine whether PDGFA exerts dose-dependent effects on alveolar fibroblasts, we cultured primary mouse or human alveolar fibroblasts in SFM supplemented with increasing concentrations of PDGFA (10, 40, or 80 ng/ml) for 5 days (Figure 3a). Immunostaining revealed a dose-dependent increase in total cell number, consistent with known mitogenic activity of PDGFA on mouse fibroblasts (Figure 3b and Extended Data Figure 4a). Accordingly, the number of Ki67^+^ cells dose-dependently increased with PDGFA, whereas ACTA2 expression was significantly suppressed at all doses relative to SFM alone (Figure 3b,c). We observed a similar pattern in primary alveolar fibroblasts derived from healthy human lungs, in which PDGFA dose-dependently increased total cell number and the number of Ki67^+^ cells, while significantly suppressing ACTA2 expression at all doses relative to SFM (Figure 3d, e and Extended Data Figure 4b).

**Figure 3:**
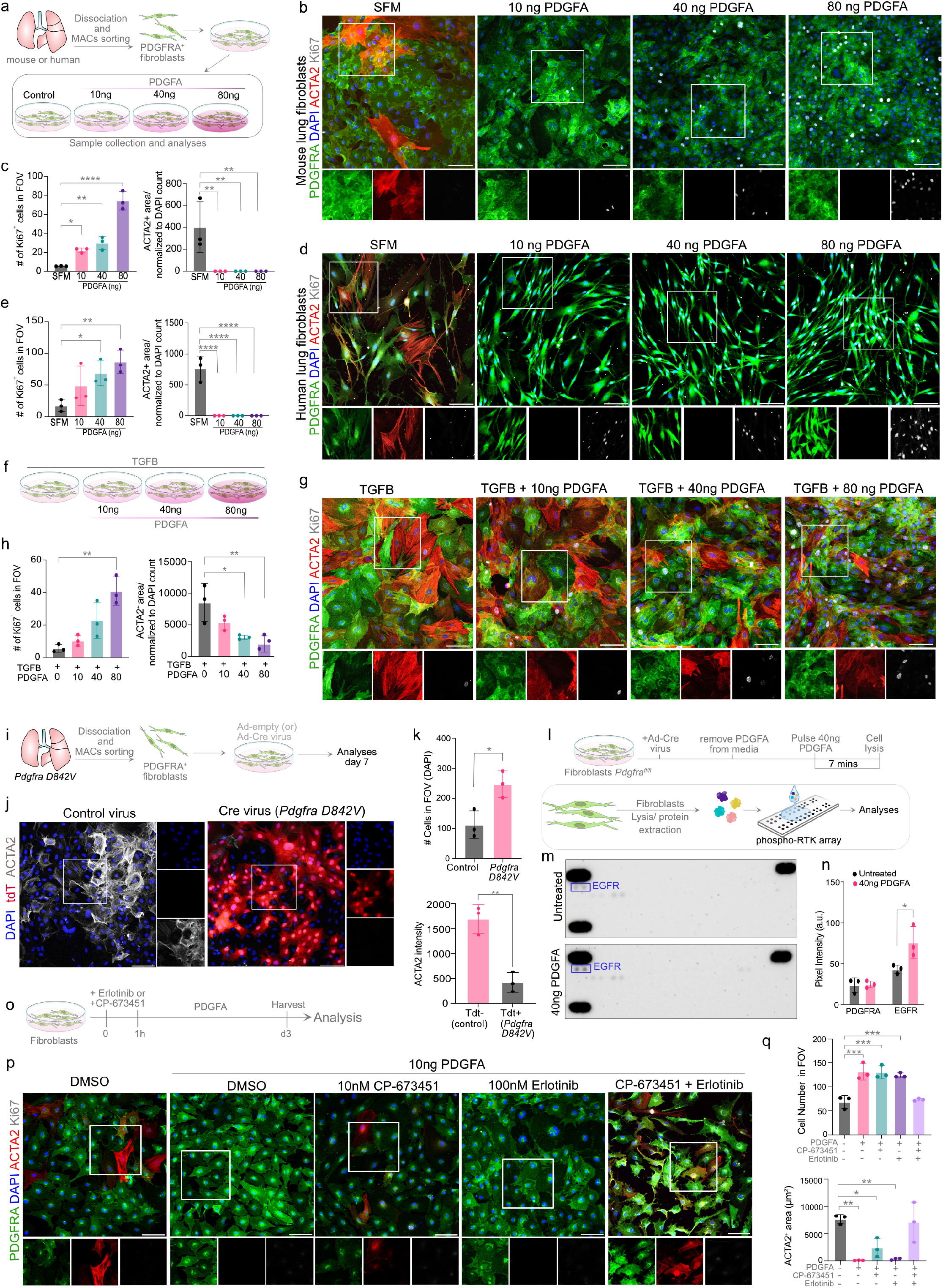
PDGFA controls alveolar fibroblast state in a dose- and context-dependent manner. a. Schematic shows timeline for alveolar fibroblast culture and PDGFA (10ng/ml, 40ng/ml, and 80ng/ml) treatments on human and mouse fibroblasts for 5 days. b. Immunostaining for PDGFRA (green), ACTA2 (red), and Ki67 (grey) in alveolar fibroblast culture, comparing SFM to increasing doses of PDGFA. Scale bar: 100μm. c. Quantification of Ki67^+^ cells in FOV and ACTA2^+^ area/normalized to DAPI count. One-way ANOVA with Dunnett’s multiple comparisons test vs. SFM; *p<0.05, **p<0.01, ****p<0.0001. n=3 mice per condition. d. Immunostaining for PDGFRA (green), ACTA2 (red), and Ki67 (grey) on human alveolar fibroblast cultures from SFM and PDGFA treatments. Scale bar: 100μm. e. Quantification of Ki67^+^ cells in FOV and ACTA2^+^ area/normalized to DAPI count. One-way ANOVA with Dunnett’s multiple comparisons test vs. SFM; *p<0.05, **p<0.01, ****p<0.0001. n=3 mice per condition. f. Schematic shows alveolar fibroblast cultures treated with PDGFA (10ng/ml, 40ng/ml, and 80ng/ml) in combination with TGFβ (10ng/ml) for 3 days. g. Immunostaining for PDGFRA (green), ACTA2 (red), and Ki67 (grey) on alveolar fibroblast cultures treated with indicated conditions. Scale bar: 100μm. h. Quantification of Ki67^+^ cells in FOV and ACTA2^+^ area/normalized to DAPI count in alveolar fibroblasts treated with indicated conditions. One-way ANOVA with Dunnett’s multiple comparisons test vs. 10ng/ml TGFβ; *p<0.05, **p<0.01. n=3 per condition. i. Schematic shows PDGFRA D842V fibroblasts culture and treatments. j. Immunostaining for tdT (red) and ACTA2 (grey) on control and *Pdgfra D842V* mutant fibroblasts. Scale bar: 100μm. k. Quantification of total cell number (upper graph) and ACTA2^+^ intensity in tdT- and tdT+ cells. Unpaired two-tailed t test; **p = 0.0032. n=3 mice per condition. l. Experimental schematic for *Pdgfra^fl/fl^* fibroblast culture, treatment with Cre virus and +/− 40ng/ml PDGFA for RTK Array analysis. m. Analysis of RTK activation in *Pdgfra^fl/fl^* fibroblasts +/− 40ng/mL PDGFA. Blue box shows EGFR phosphorylation spots. n. Quantification of EGFR phosphorylation. Two-way ANOVA with Sidak’s multiple comparisons test; *p=0.0139. n=3 per condition. o. Schematic shows experimental design for fibroblast culture and treatment with PDGFRA inhibitor CP-673451 and EGFR inhibitor Erlotinib. p. Immunostaining for PDGFRA (green), ACTA2 (red), and Ki67 (grey) in alveolar fibroblast cultures after 3-days of CP-673451 (10 nM), Erlotinib (100 nM), or in combination and +/− 10ng/ml PDGFA treatment in SFM. Scale bar: 100μm. q. Quantification of total cell number in FOV and ACTA2^+^ area (µm²). One-way ANOVA with Dunnett’s multiple comparisons test vs. SFM; *p<0.05, **p<0.01, ***p<0.001. n=3 mice per condition. DAPI stains nuclei in all panels. Statistical significance is indicated by asterics.In all graphs data are presented as mean ± SD.

We next asked whether PDGFA modulates fibroblast activation in the presence of an active pro-fibrotic signal. To this end, we cultured alveolar fibroblasts with TGFβ (10 ng/ml) alone or in combination with increasing concentrations of PDGFA (10, 40, or 80 ng/ml) for 3-days (Figure 3f). Immunostaining for PDGFRA, ACTA2, and Ki67 showed that the combination of TGFβ and PDGFA dose-dependently increased both total cell number and the number of Ki67^+^ cells relative to TGFβ alone (Figure 3g, h and Extended Data Figure 4c). Notably, higher concentrations of PDGFA in the presence of TGFβ showed a trend toward reduced ACTA2 expression relative to TGFβ alone, suggesting that PDGFA partially tempers TGFβ-induced activation while still permitting myofibroblast differentiation (Figure 3g,h). By contrast, a lower dose of PDGFA (10 ng/ml) appeared to promote expansion of ACTA2^+^ myofibroblasts when combined with TGFβ, increasing the number of Ki67^+^ cells while maintaining their ACTA2^+^ state (Figure 3h). Together, these findings demonstrate that PDGFA acts in a context- and dose-dependent manner. Overall, these data support a model in which PDGFA, contrary to the prevailing view of its uniformly pro-fibrotic activity, can suppress acquisition of the myofibroblast fate, with its effects shaped by both dose and the presence of additional signals such as TGFβ.

### PDGFRA is sufficient for PDGFA signaling but is functionally compensated by EGFR in alveolar fibroblasts

Having established that PDGFA treatment is sufficient to maintain alveolar fibroblast identity, we next asked whether activation of PDGFRA is sufficient to reproduce these effects. We isolated alveolar fibroblasts from *Pdgfra-D842V;fsfTdt* mice (hereafter *Pdgfra-D842V*), in which Cre recombination concomitantly activates a constitutively active PDGFRA mutant together with tdTomato expression, allowing recombined tdT^+^ cells to be distinguished from non-recombined tdT^-^ cells within the same culture (Figure 3i). Fibroblasts were cultured under serum-free conditions, infected with control adenovirus or adenovirus expressing Cre recombinase (AdV-Cre), and analyzed after 7 days. Activation of *PDGFRA D842V* significantly increased cell number while reducing ACTA2 expression relative to controls (Figure 3j,k). Recombination efficiency was confirmed by quantifying the percentage of tdT^+^ cells per field of view (Extended Data Figure 4d). Within the same cultures, tdT^+^ cells exhibited reduced ACTA2 expression and ACTA2 coverage compared with neighboring tdT^-^ cells (Extended Data Figure 4e-g), demonstrating a cell-intrinsic effect of PDGFRA activation. Importantly, these phenotypes occurred in the absence of exogenous PDGFA, indicating that constitutive PDGFRA activation phenocopies PDGFA treatment independently of ligand availability. Together, these findings demonstrate that PDGFRA activation is sufficient to suppress myofibroblast differentiation while promoting fibroblast proliferation.

We next asked whether PDGFRA is required for these effects. Alveolar fibroblasts isolated from *Pdgfra^fl/fl^* mice were cultured in SFM and treated with AdV-Cre to delete *Pdgfra*, followed by culture in the presence or absence of PDGFA (Extended Data Figure 4h). Unexpectedly, loss of *Pdgfra* did not significantly alter fibroblast cell number or ACTA2 expression compared with control fibroblasts (Extended Data Figure 4i,j). This was observed in SFM alone, SFM supplemented with PDGFA, and across different concentrations of TGFβ (Extended Data Figure 4i,j). Thus, although PDGFRA activation is sufficient to reproduce the effects of PDGFA, PDGFRA is dispensable for fibroblasts to respond to PDGFA, suggesting the existence of a compensatory signaling mechanism.

To identify signaling pathways that might compensate for loss of PDGFRA, we profiled receptor tyrosine kinase (RTK) activity using a phospho-RTK array. Alveolar fibroblasts isolated from *Pdgfra*^fl/fl^ mice were treated with AdV-Cre to delete *Pdgfra* and subsequently stimulated with PDGFA (40 ng/ml). Phospho-RTK profiling revealed increased EGFR phosphorylation following PDGFA treatment in *Pdgfra*-deficient fibroblasts compared with untreated controls (Figure 3l-n), identifying EGFR as a candidate compensatory receptor. To functionally test this possibility, primary alveolar fibroblasts were treated with CP-673451 or erlotinib to inhibit PDGFRA or EGFR, respectively, in the presence or absence of PDGFA (Figure 3o). Immunostaining for ACTA2, PDGFRA, and Ki67 revealed that inhibition of either PDGFRA or EGFR alone was insufficient to block PDGFA-induced proliferation or suppression of ACTA2. In contrast, combined inhibition of PDGFRA and EGFR abolished these PDGFA-induced phenotypes (Figure 3p,q). Higher concentrations of CP-673451 alone also blocked the effects of PDGFA, likely reflecting broad range inhibitor activity at higher doses (Extended Data Figure 4k-m). Together, these findings demonstrate that PDGFRA activation is sufficient to reproduce the effects of PDGFA on alveolar fibroblasts but is dispensable and identify EGFR as a compensatory receptor that sustains PDGFA responses when PDGFRA activity is impaired.

### Phosphoproteomics identifies SHP2-associated signaling downstream of PDGFA in alveolar fibroblasts

Having established that PDGFA maintains alveolar fibroblast identity, we next asked which downstream signaling programs regulate fibroblast state. Purified alveolar fibroblasts were stimulated with PDGFA at 80 ng/ml, TGFβ at 20 ng/ml, or a combination of PDGFA (80 ng/ml) and TGFβ (20 ng/ml), hereafter referred to as Combo, followed by phosphoproteomic analysis. Samples were collected 7 minutes after PDGFA treatment and 30 minutes after TGFβ treatment to capture acute signaling responses (Figure 4a). Following batch normalization and quality-control assessment, we quantified differentially abundant phosphorylation sites. Relative to SFM controls, differential phosphosite analysis identified 1,666 significantly altered sites in TGFβ-treated cells (1,345 increased and 321 decreased), 1,836 in PDGFA-treated cells (1,593 increased and 243 decreased), and 2,176 in Combo-treated cells (1,727 increased and 449 decreased) (Extended Data Figure 5a and and Supplementary Data Table 1).

**Figure 4:**
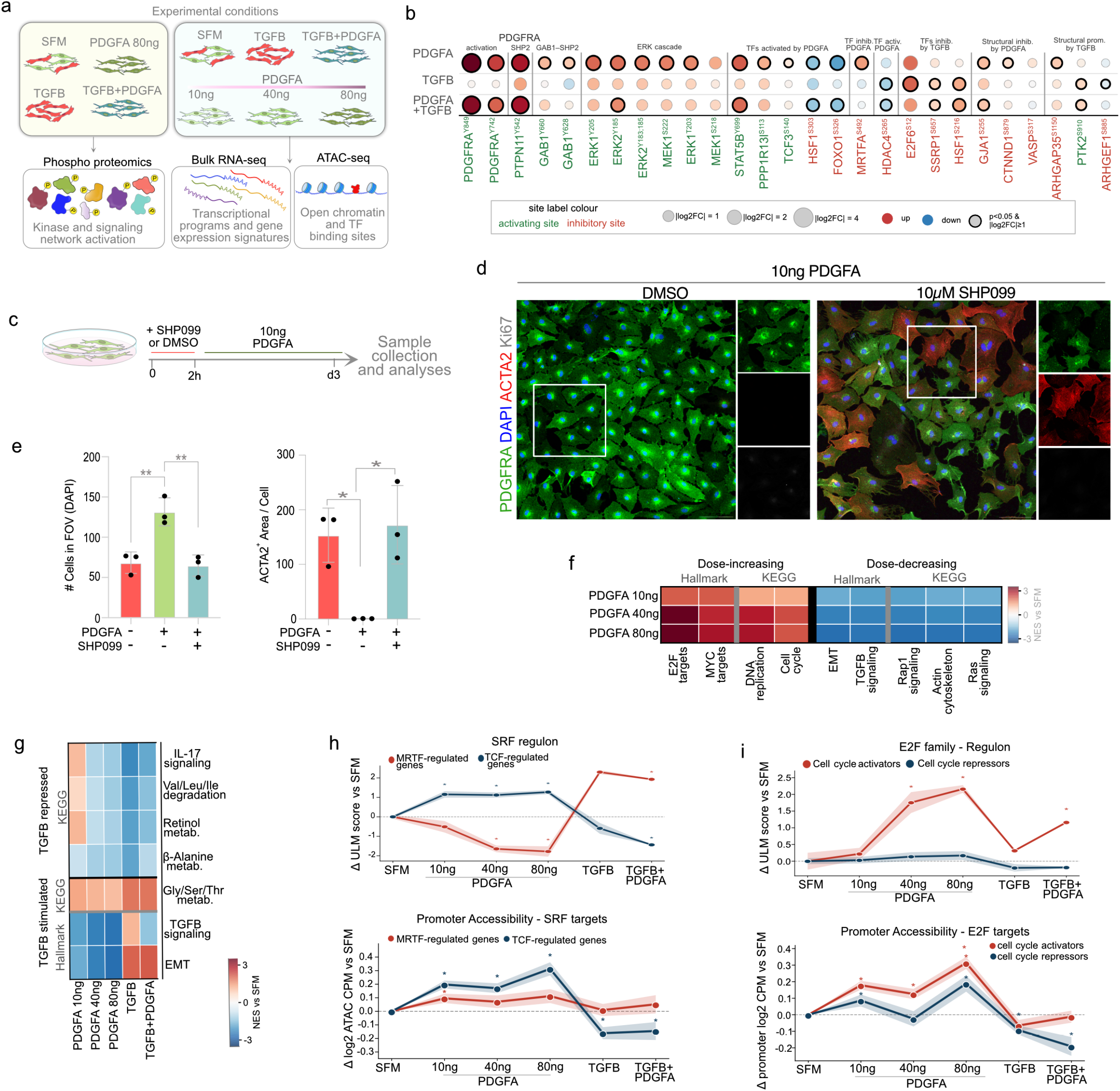
Multi-omic profiling reveals transcriptional and signaling programs downstream of PDGFA. a. Schematic showing experimental design for alveolar fibroblasts treated with SFM, PDGFA (10– 80 ng/ml), TGFβ, or TGFβ+PDGFA followed by phosphoproteomics, bulk RNA-seq, and ATAC-seq. b. Phosphosite changes with PDGFA, TGFβ, or TGFβ+PDGFA compared to SFM, grouped by signaling node. Label color denotes activating (green) or inhibitory (red) site. c. Schematic shows timeline for alveolar fibroblast culture and treatment with SHP099 (10 µM) or DMSO pretreatment 2h before 10 ng/ml PDGFA, harvested 3-days after treatment. d. Immunostaining for PDGFRA (green), ACTA2 (red), and Ki67 (gray) in DMSO- vs. SHP099-treated cells with 10 ng/ml PDGFA. Scale bar: 100μm. e. Quantification of total cell number in FOV (left). One-way ANOVA with Tukey’s multiple comparisons test (all pairwise comparisons); **p <0.01. n=3 per condition. Quantification of ACTA2^+^ area/normalized to cell number in FOV (right). One-way ANOVA with Tukey’s multiple comparisons test (all pairwise comparisons); *p<0.05. n=3 per condition. f. Hallmark/KEGG pathway enrichment (NES) increasing or decreasing with PDGFA dose vs. SFM (GSEA). g. Hallmark/KEGG pathway enrichment (NES) for TGFβ-stimulated or -repressed pathways across PDGFA dose, TGFβ, and TGFβ+PDGFA vs. SFM. h. SRF regulon activity for MRTF- (red) and TCF-regulated (blue) genes across conditions. Promoter accessibility (Δlog2 ATAC CPM vs. SFM, TSS ±5 kb) for SRF targets. Shaded area represents ±SEM; *p<0.05 vs. SFM. i. E2F regulon activity for cell-cycle activators (red) and repressors (blue), as in (h). Promoter accessibility (Δlog2 ATAC CPM vs. SFM, TSS ±5 kb) for E2F targets. Shaded area represents±SEM; *p<0.05 vs. SFM. DAPI stains nuclei. Statistical significance is indicated by asterics.

We next examined the signaling proteins and pathways associated with these phosphosite changes. PDGFA and Combo treatment increased phosphorylation of proteins associated with receptor and proliferative signaling, including PDGFRA^Y849^, PTPN11^Y542^, SPRY2^T60^, MAPK1^Y185^, and AKT1^T479^. In contrast, TGFβ and Combo treatment increased phosphorylation of proteins associated with cytoskeletal regulation, including ARPC1B^S310^, ZEB2^T994^, LIMK1^S261^, and VIM^S49^ (Figure 4b and Extended Data Figure 5b). Consistent with previous studies, pathway analysis of TGFβ-responsive phosphosites showed predominant enrichment of PLC/PKC/calcium signaling, receptor tyrosine kinase signaling, and PI3K/AKT/mTOR pathways ^30,31^. PDGFA-responsive phosphosites were enriched for expected pathways, including RAS/MAPK/ERK signaling and adaptor protein signaling. Significantly, STAT5B^Y699^ activation is increased in PDGFA and Combo treatments, as well as increases in activating phosphosites for TCF3^S140^ and PPP1R13l^S113^, and a decrease in inhibitory sites for FOXO1^S326^ and HSF1^S303^. PDGFA treatment also uniquely inhibits the transcription factor MRTFA^S492^. TGFβ treatment also impacted select transcription factors within 30 minutes of treatment. E2F6^S12^, HSF1^S216^ and SSRP1^S657^ inhibitory sites were all increased in TGFβ treatment (Figure 4b). In addition to transcription factors, PDGFA also negatively regulates cytoskeletal proteins VASP^S317^, GJA1^S255^, and CTNND1^S879^. TGFβ on the other hand, promotes cytoskeletal proteins PTK2^S910^ and ARHGEF1^S885^ (Figure 4b).

Kinase activity inference using benchmarKIN and phosphosite annotation using SIGNOR further supported activation of an SHP2-ERK signaling axis downstream of PDGFA. Increased phosphorylation of SHP2-Y542 was accompanied by phosphorylation of GAB1^Y628^ and GAB1^Y660^, sites associated with SHP2-GAB1 complex formation, as well as MEK1^S218/S222^, ERK1^T203/Y205^, and ERK2^Y185^. Together, these phosphorylation events are consistent with engagement of an SHP2-associated ERK signaling cascade downstream of PDGFA (Figure 4b and Extended Data Figure 5a).

To determine whether SHP2 activity contributes functionally to PDGFA-mediated regulation of fibroblast state, we cultured alveolar fibroblasts in SFM and pretreated them with the SHP2 inhibitor SHP099 for 2h before addition of PDGFA (10 ng/ml) (Figure 4c). Immunostaining for PDGFRA, ACTA2, and Ki67 showed that SHP099 co-treatment blocked both the PDGFA-induced increase in cell number and the reduction in ACTA2 expression (Figure 4d,e). Together, these findings identify SHP2 activity as a key mediator of PDGFA-induced fibroblast proliferation and suppression of myofibroblast differentiation.

### Transcriptome and chromatin accessibility profiling define dose- and context-dependent programs associated with fibroblast state

We next asked which transcriptional and chromatin programs accompany the dose- and context-dependent effects of PDGFA (10, 40, or 80 ng/ml) in comparison with TGFβ (10ng/ml) and Combo (PDGFA (80ng/ml) plus TGFβ (10ng/ml)) treatments. We profiled and analyzed bulk RNA-seq and ATAC-seq data collected 72h after treatment (Figure 4a and Supplementary Data Table 2). Differential gene expression analysis using DESeq2 identified gradual changes with increasing PDGFA concentration. These include downreguation of *Acta2*, *Tagln*, *Fn1*, and *Bmp4*, whereas upregulated genes include *Spon2*, *Sparcl1*, and *Timp1* (Extended Data Figure 5c). MSigDB Hallmark analysis of genes whose expression increased with PDGFA dose identified enrichment of gene sets related to cell cyce, DNA replication, E2F and MYC targets (Figure 4f). Dose-decreasing genes were enriched for epithelial-mesenchymal transition (EMT), actin cytoskeleton, TGFβ, and Rap1signaling (Figure 4f). Together, these transcriptional changes were consistent with the preceding immunostaining results, in which increasing PDGFA concentrations promoted fibroblast proliferation while suppressing ACTA2 expression. Whereas TGFβ treatment showed enrichment of EMT, and TGFβ signaling gene sets. (Figure 4g). Gly/Ser/Thr metabolism was induced by both PDGFA and TGFβ, although more weakly by PDGFA. Other TGFβ-repressed pathways showed dose-dependent responses to PDGFA, with IL-17 signaling and several metabolic pathways enriched at low PDGFA but suppressed at higher doses. Combination treatment showed mixed effects, maintaining the EMT signature while reducing TGFβ signaling compared with TGFβ alone.

ATAC-seq data analysis revealed enrichment of chromatin loci corresponding to genes associated with PDGFA, TGFβ, and Combo treatments. Specifically, chromVAR analysis revealed increased chromatin accessibility in genomic regions with predicted motifs for EGR2, PRRX1, TCF21, FOSL2, and MEIS2 in PDGFA treatments (Extended Data Figure 5d and Supplementary Data Table 3). TGFβ treated cells showed enrichment of genomic regions with motifs for SMAD3/5, GATA1, and ZNF317 (Extended Data Figure 5d). Notably, Combo-treated samples showed enrichment of motifs shared with PDGFA or TGFβ conditions including SMAD3/5, JUN, FOSL2, TBXT and TP53 (Extended Data Figure 5d). Together, transcriptome and chromatin accessibility profiles revealed unique and shared transcriptional features in PDGFA, TGFβ, and Combo treatments.

### Multi-omic integration links PDGFA responses to divergent SRF/MRTFA and E2F regulatory modules to antagonize TGFβsignaling

We next integrated the phosphoproteomic, transcriptomic, and chromatin accessibility datasets to determine how upstream signaling is coupled to transcriptional programs that regulate fibroblast proliferation and myofibroblast differentiation. Among the phosphosites altered by PDGFA treatment, MRTFA^S492^, an inhibitory phosphorylation site, was significantly enriched (Figure 4b). Given the established role of MRTFA as a cofactor for serum response factor (SRF), we asked whether PDGFA differentially regulates SRF-dependent transcriptional programs associated with myofibroblast differentiation and proliferation ^32^. Stratification of SRF target genes revealed two divergent programs: an MRTFA-associated module enriched for myogenic and myofibroblast-associated genes, and a ternary complex factor (TCF)-associated module comprising non-myogenic SRF targets (Figure 4h and Extended Data Figure 5e).

Expression of TCF-associated targets increased at 10 ng/ml PDGFA and remained elevated at higher concentrations, whereas expression of MRTFA-associated targets decreased with PDGFA treatment. TGFβ shifted this balance toward the MRTFA-associated program, which remained elevated under Combo treatment (Figure 4h). Promoter accessibility within the TCF-associated regulon broadly paralleled its transcriptional response. In contrast, chromatin accessibility within the MRTFA-associated regulon remained largely unchanged relative to SFM despite substantial changes in target-gene expression (Figure 4h). Thus, suppression of MRTFA-associated transcription was not accompanied by a corresponding reduction in promoter accessibility. Together with increased phosphorylation of MRTFA-S492, these findings suggest that PDGFA suppresses the myofibroblast-associated SRF program primarily through regulation of MRTFA activity rather than through widespread changes in chromatin accessibility.

In parallel, analysis of shared E2F family targets identified enrichment of E2F-associated programs under PDGFA-treated conditions. We further resolved these targets into cell-cycle and tumor-suppressor modules. Expression of E2F cell-cycle targets increased modestly at 10 ng/ml PDGFA and more strongly at higher doses. TGFβ markedly suppressed this program, whereas Combo treatment partially restored cell-cycle target expression toward levels observed with PDGFA alone. By contrast, expression of the E2F tumor-suppressor module remained largely unchanged across conditions (Figure 4i and Extended Data Figure 5f,g), indicating that the PDGFA-responsive E2F program is preferentially associated with cell-cycle control. Consistent with this transcriptional response, ATAC-seq analysis revealed increased accessibility at regulatory regions associated with both E2F modules following PDGFA treatment, with a more pronounced increase at cell-cycle-associated loci (Figure 4i). TGFβ reduced accessibility across both modules, whereas Combo treatment partially restored accessibility at cell-cycle-associated regions while further reducing accessibility associated with the tumor-suppressor module (Figure 4i). Thus, chromatin accessibility and transcriptional output were concordant for the proliferative E2F program, whereas changes in accessibility within the tumor-suppressor module were not accompanied by comparable changes in gene expression.

Together, these multi-omic analyses support a model in which PDGFA simultaneously promotes an E2F-associated proliferative program while suppressing MRTFA-associated SRF transcription linked to myofibroblast differentiation. In the presence of TGFβ, PDGFA partially restores the E2F cell-cycle program without reversing the predominance of MRTFA-associated transcription. This regulatory mechanism is consistent with our functional data showing that PDGFA promotes fibroblast proliferation and maintains their identity while antagonizing myofibroblast differentiation.

### PDGFA treatment reverses human fibrotic lung myofibroblasts into fibroblasts ex vivo

Our findings above indicate that PDGFA antagonizes TGFβ-driven fibroblast activation by suppressing myofibroblast-associated programs. Consistent with this, high-dose PDGFA (80 ng/ml) reduced ACTA2^+^ area even in the presence of TGFβ (Figure 3g, h). We therefore asked whether PDGFA could reverse an already established myofibroblast state rather than simply prevent fibroblast activation. To test this, we first induced myofibroblast differentiation in mouse alveolar fibroblasts with TGFβ (10 ng/ml), then switched the cultures to SFM with or without PDGFA (10 or 80 ng/ml) for 4 days (Figure 5a). Fibroblast state was assessed by immunostaining for PDGFRA, ACTA2, and Ki67 (Figure 5b). By 2-days after PDGFA treatment, fibroblasts had begun to lose ACTA2 expression and showed an increase in total cell number (Extended Data Figure 6a-c). By day-4, PDGFA treatment produced a dose-dependent increase in total cell number together with a reduction in ACTA2 expression relative to SFM alone (Figure 5c), indicating that PDGFA can promote reversion of an established myofibroblast state.

**Figure 5:**
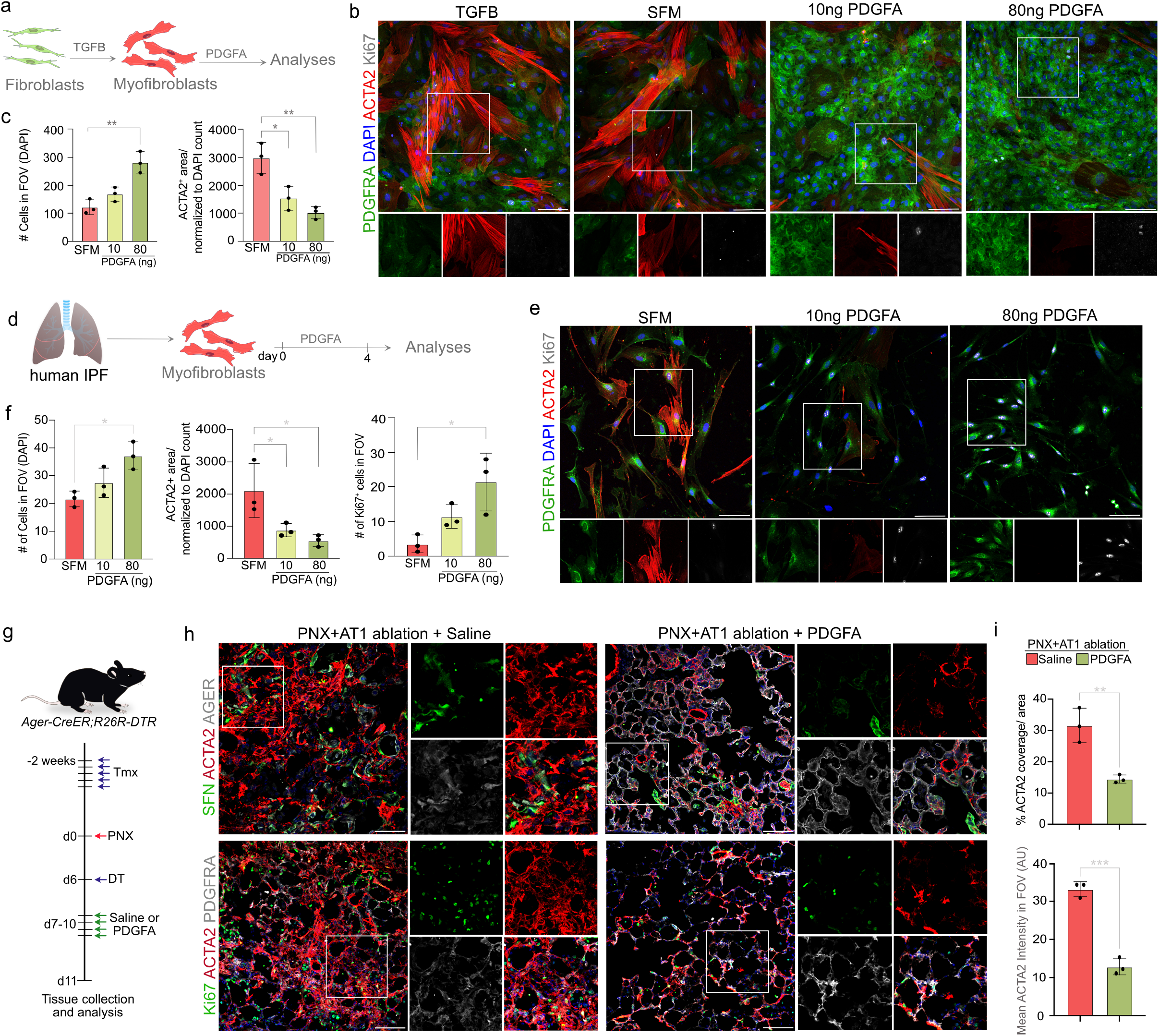
PDGFA promotes myofibroblast reversion and reduces fibrosis *in vivo*. a. Schematic shows treatement regimen for PDGFA and TGFβ on primary mouse alveolar fibroblasts. b. Immunostaining for PDGFRA (green), ACTA2 (red), and Ki67 (grey) in myofibroblasts treated with SFM, 10 ng/ml PDGFA, or 80 ng/ml PDGFA for 4 days. Scale bar: 100 µm. c. Quantification of total cell number in FOV and ACTA2^+^ area/normalized to DAPI count. One-way ANOVA with Dunnett’s multiple comparisons test vs. SFM; *p<0.05, **p<0.01. n=3 mice per condition. d. Schematic shows shows treatement regimen for PDGFA on myofibroblasts from human IPF lungs. e. Immunostaining for PDGFRA (green), ACTA2 (red), and Ki67 (grey) on human IPF myofibroblasts treated with SFM, 10 ng/ml PDGFA, or 80 ng/ml PDGFA for 4-days. Scale bar: 100 µm. f. Quantification of total cell number, Ki67^+^ cells in FOV and ACTA2^+^ area/normalized to DAPI count. One-way ANOVA with Dunnett’s multiple comparisons test vs. SFM; *p<0.05. n=3 biological replicates per condition. g. Schematic shows timeline for *in vivo* PDGFA adminsitrtaion in a progressive fibrosis model. h. Immunostaining for SFN (green), ACTA2 (red), AGER (grey) (top), and Ki67 (green), ACTA2 (red) and PDGFRA (grey) (bottom) on lung tissues from saline- and PDGFA-treated *AgercreER;R26R-DTR* mice (n=3 per group). Scale bar: 100μm. i. Quantification of ACTA2 coverage and mean ACTA2 intensity in FOV (AU). Unpaired two-tailed t-test; **p=0.0073, ***p=0.0003. n=3 mice per condition. DAPI stains nuclei in all panels. Statistical significance is indicated by asterics. In all graphs data are presented as mean ± SD.

To determine whether this effect extends to disease-relevant human cells, we isolated myofibroblasts from idiopathic pulmonary fibrosis (IPF) lungs, cultured them in SFM, and treated them with PDGFA (Figure 5d). Immunostaining for PDGFRA, ACTA2, and Ki67 showed that PDGFA similarly increased the number of human IPF-derived fibroblasts in a dose-dependent manner while reducing ACTA2 expression (Figure 5e,f). Consistent with this increase in cell number, Ki67 immunostaining revealed a significant increase in proliferating cells following PDGFA treatment (Figure 5f). Together, these findings demonstrate that PDGFA promotes reversion of an established myofibroblast state in both mouse alveolar fibroblasts and human IPF-derived myofibroblasts. These data further suggest that restoring PDGFA signaling may be sufficient to shift activated fibroblasts away from a myofibroblast state even in a disease-relevant context.

### PDGFA treatment attenuates fibrosis in vivo

To determine whether PDGFA could reduce fibrotic remodeling *in vivo*, we used a previously described progressive fibrosis model based on sequential injury, consisting of pneumonectomy (PNX) followed by AT1 cell ablation ^21^. *Ager-creER;R26R-DTR* mice received tamoxifen 2 weeks before PNX, followed by diphtheria toxin on day 6 to ablate AT1s. Mice then received saline or PDGFA on days 7-10, and lungs were collected on day 11 (Figure 5g). Immunostaining for SFN, AGER, ACTA2, Ki67, and PDGFRA revealed that saline-treated lungs contained dense ACTA2^+^ fibrotic regions, whereas PDGFA-treated lungs showed a thinner interstitium and reduced ACTA2 expression (Figure 5h). Similarly, immunostaining for SPC, ACTA2, and LEL showed that saline-treated lungs exhibited disorganized alveolar architecture with prominent ACTA2^+^ fibrotic regions, whereas PDGFA-treated lungs displayed better-preserved alveolar architecture with thinner ACTA2^+^ interstitial regions (Extended Data Figure 6d). Quantification confirmed that PDGFA-treated lungs had significantly reduced ACTA2 expression compared with saline-treated controls (Figure 5i and Extended Data Figure 6e), together with reduced tissue area and reduced percent tissue coverage (Extended Data Figure 6e).

Together, these findings demonstrate that PDGFA treatment attenuates fibrotic remodeling in vivo in a progressive lung fibrosis model. These data further support AT1-derived PDGFA signaling as a potential therapeutic axis for preserving fibroblast identity and limiting fibrotic remodeling in the injured lung.

## Discussion

Our study identifies AT1-derived PDGFA as a tonic homeostatic signal that communicates epithelial state to alveolar fibroblasts and maintains their identity under steady-state conditions. Loss of AT1s or depletion of AT1-derived PDGFA was sufficient to activate alveolar fibroblasts and initiate an injury-like tissue response without an exogenous insult. Conversely, restoration of PDGFA signaling suppressed fibroblast activation, promoted transition away from established myofibroblast states, and attenuated fibrotic remodeling in vivo. These findings support a model in which the AT1-PDGFA-fibroblast axis functions as a homeostatic checkpoint linking epithelial integrity to the initiation and resolution of tissue repair.

This model suggests that cellular quiescence requires continuous instructive communication between tissue compartments. AT1s occupy an extensive interface with the alveolar interstitium and are therefore well positioned to regulate neighboring stromal cells through trophic signaling. Loss of AT1s could reduce local PDGFA availability, releasing fibroblasts from homeostatic restraint and permitting their transition into injury-associated states. Further, our data from repeat AT1 ablation of extensive loss of PDGFA led to alveolar simplification, suggesting that dose of PDGFA and AT1 cell numbers dictate the outcomes. Conversely, our observation that *Pdgfa* expression progressively increases during AT2-to-AT1 differentiation suggests that epithelial regeneration could restore this restraint. Thus, AT1s may function as tissue organizers whose loss and regeneration coordinate fibroblast activation and return toward homeostasis. Previous work demonstrated that epithelial SHH signaling actively maintains quiescence of the adjacent peribronchiolar mesenchyme, with attenuation of this signal after injury permitting mesenchymal cell proliferation and its restoration contributing to resolution ^10^. Although previous studies have described a critical role for PDGFA and PDGFRA in lung development, our findings reveal a new paradigm in distal alveolar niche, where loss of AT1s or AT1-derived PDGFA is sufficient to release alveolar fibroblasts from quiescence and trigger a broader cascade of injury-like responses ^33–36^.

Our findings also challenge the prevailing view that PDGF signaling promotes fibroblast activation and fibrosis ^27,37,38^. Instead, the effects of PDGFA depend on dose, cellular context, and fibroblast state. At relatively low concentrations, PDGFA maintained alveolar fibroblast identity with limited proliferation. Increasing concentrations promoted proliferation while continuing to suppress ACTA2 expression, indicating that proliferation and myofibroblast differentiation are separable outputs. High-dose PDGFA also reduced ACTA2 expression following TGFβ-induced differentiation and in fibroblasts isolated from human IPF lungs. Thus, PDGFA can expand fibroblast populations while restraining acquisition or maintenance of the contractile myofibroblast state. This is in line with previous studies that showed fibroproliferation without scarring upon exporession of constitutive PDGFRA signaling ^38^.

Our multi-omic analyses provide a mechanistic framework for this separation. Acute PDGFA stimulation engaged PDGFRA-, SHP2-, and ERK-associated signaling, while SHP2 inhibition blocked the proliferative and ACTA2-suppressive effects of PDGFA. This is in line with previous reports that SHP2 represses fibroblast to myofibroblasts ^39^. PDGFA increased phosphorylation of MRTFA at an inhibitory site and suppressed MRTFA-associated SRF targets while favoring TCF-associated programs. In parallel, E2F-associated cell-cycle transcription increased with PDGFA dose. TGFβ favored MRTFA-associated transcription and suppressed the E2F program. Combined treatment partially restored E2F-associated transcription without reversing the predominance of the MRTFA-associated program, consistent with fibroblasts proliferating while retaining myofibroblast features. A similar scenario has been described in fibroblast to myofibroblasts conversion during wound healing ^40^. These findings suggest that the relative activities of PDGFA- and TGFβ-responsive programs help determine fibroblast state.

This balance may coordinate fibroblast dynamics during injury and repair. Reduced PDGFA availability following AT1 loss, together with increased TGFβ and other injury-associated signals, could favor myofibroblast differentiation. Subsequent restoration of AT1-derived PDGFA could shift fibroblasts toward a homeostatic state. The ability of PDGFA to reduce myofibroblast features in mouse and human IPF-derived fibroblasts and attenuate remodeling in a progressive fibrosis model raises the possibility of therapeutically promoting this transition. Such an approach could preserve fibroblasts needed for alveolar structure and repair, although its effects would likely depend on dose, the surrounding signaling environment, and the state of the responding cells.

Several questions remain. The spatial and temporal regulation of PDGFA availability during injury and the responsiveness of distinct fibroblast states require further study. EGFR compensation following loss of PDGFRA signaling also suggests that receptor context influences these responses. Moreover, whether PDGFA-treated myofibroblasts fully reacquire the molecular and functional properties of homeostatic alveolar fibroblasts remains unresolved. Together, our findings identify the AT1-PDGFA-fibroblast axis as a homeostatic circuit linking epithelial integrity to stromal state and suggest that disruption and restoration of tonic epithelial signals can coordinate the initiation and resolution of tissue repair.

## Methods

### Animals

*Sftpc-cre/ERT2* (Sftpc-CreER) (028054, Jackson Laboratory ^41^), *Rosa26R-lsl-DTA* (006331, Jackson Laboratory ^42^), *Rosa26R-lsl-DTR* (007900, Jackson Laboratory ^43^), *Pdgfra-creERT2* (032770, Jackson Laboratory ^44^), *PdgfraK.Flp;fSf-Tomato* (038627, Jackson Laboratory ^45^), *Pdgfra^fl/fl^* (006492, Jackson Laboratory ^46,47^), *Ager cre/ERT2* (036942, Jackson Laboratory ^44^), *Pdgfa^fl/fl^* ^48^ crossed with Ager-CreER to achieve AT1-specific conditional deletion of *Pdgfa*, *Cdh5-creER* (006137, Jackson Laboratory ^49^), CD169-DTR (kindly provided by the Dee Gunn laboratory, Duke University ^50^), and C57BL/6J (stock number 000664, Jackson Laboratory). Mice were maintained on a C57BL/6J background. For all loss of function and ablation experiments, 3–5 doses of tamoxifen (0.2 mg/g body weight) (Sigma-Aldrich) were administered intraperitoneally, except for AT1 ablation, for which five doses of tamoxifen (0.4 mg/g body weight) were administered. DT was administered at a dose of 3 μg/mouse (single ablation) or 3 μg/mouse + 1 μg/mouse (repeat ablation) via IP administration. Both male and female mice between 8 and 16 weeks of age were used for experiments. All animal experiments were approved by the Duke University Institutional Animal Care and Use Committee in accordance with US National Institutes of Health guidelines.

### Partial left lobe pneumonectomy

Partial left-lobe PNX was performed as previously described ^51^. Briefly, animals were anesthetized by inhalation of 3% isoflurane in an induction chamber, and anesthesia was maintained via peroral tracheal intubation with a 22G catheter (ventilation set with MiniVent, 200-300ul/stroke). The left thoracic wall was shaved and disinfected with betadine and ethanol. Buprenorphine-SR was administered subcutaneously as an analgesic. A 1-1.5cm incision was made over the left rib cage and skin retracted, followed by a small incision in the intercostal space between the fifth and sixth ribs and expanded using a retractor. The left lobe of the lung was lifted out of the rib cage with forceps, and the pulmonary vessels and left bronchi were ligated with a surgical clip. The left lung was resected, and the rib cage sutured shut. Outer skin was stapled shut. Bupivicaine (50μL) was administered locally to the incision site. Mechanical ventilation was ceased once mice returned to spontaneous breathing on room air. Mice were then extubated and allowed to recover in a cage on a heat source. Saline, 1ml at 0.9% sodium chloride (USP), was given intraperitoneally to replace fluid/blood loss. Animals were monitored every 15 min until ambulatory and then at least twice daily for 72h. For double injury, progressive fibrosis (PNX + AT1 ablation) model, *Ager-CreER* mice were administered five doses of tamoxifen (0.4 mg/g body weight) and PNX performed as above. DT (3μg/mouse) was intraperitoneally administered on day-6 and PDGFA was intranasally administered under isoflurane anesthesia at a dose of 2μg in normal saline.

### Tissue preparation and sectioning

Lungs were perfused with PBS to remove blood, then inflated with 4% Paraformaldehyde (PFA) and incubated for 4–6h or overnight at 4°C. Lung lobes were separated, washed in PBS, and incubated overnight in 30% sucrose at 4°C. Lobes were subsequently incubated in 1:1 30% sucrose:OCT for 1h followed by embedding in OCT blocks and cryosectioning. For all tissue imaging, sections were collected at 8μm.

### Mouse lung tissue dissociation and cell isolation

Lung dissociation was performed as described previously ^21,24^. Briefly, lungs were inflated with an enzyme dissociation solution (450U/ml Collagenase I (Worthington, LS004197), 5U/ml *Dispase* (Corning, 354235), and 0.33U/ml DNase I (Roche, 10104159001), in DMEM). The lung lobes were separated, minced, and incubated in the enzyme solution at 37°C for 25–35 min. Dissociation was quenched with an equivalent volume 10% FBS/DMEM and strained through a 100μm strainer. The cell pellet was resuspended in red blood cell lysis buffer (100 μM EDTA, 10 mM KHCO_3_, 155 mM NH_4_Cl) for 2 min, followed by quenching with 10% FBS/DMEM and filtration through a 40μm strainer.

### Mouse Fibroblasts purification, culture, and treatment

Magnet assisted cell sorting was performed as described previously ^52^. Briefly, the dissociated cells were resuspended in 1mL of MACs buffer (PBS, pH 7.2, 0.5% bovine serum albumin (BSA), and 2 mM EDTA). MACs antibodies used for sorting include the following: CD45 MicroBeads, mouse (Milenyi Biotec, 130-052-301, 1:100), CD31 MicroBeads, mouse (Miltenyi Biotec, 130-097-418, 1:100), EpCAM MicroBeads, mouse (130-105-958, 1:100), LSEC MicroBeads, mouse (Miltenyi Biotec, 130-092-007, 1:100). Fibroblasts were cultured in SFM (Serum Free Media) containing; DMEM/F12 (Gibco, 12634-010) with Antibiotic/Antimycotic (Gibco 15240062), HEPES (Thermo Fisher Scientific, 15630080). GlutaMAX (Thermo Fisher Scientific, 35050061), B27 supplement (Thermo Fisher Scientific, 17504044), N2 supplement (Thermo Fisher Scientific, 17502048), N-acetylcysteine (Sigma-Aldrich, A9165). After isolation, cells were resuspended in media and seeded on 5% matrigel coated plates (Corning, 354230). Briefly, Matrigel was serially diluted in DMEM/F12 to a concentration of 5%, followed by well-coating at 37°C for 30 min. Next, Matrigel was removed and fibroblasts were seeded onto the plates. Media was changed every 2– 3 days. PDGFA was added to the culture media at 10ng/ml, unless otherwise stated (BioLegend, 776306). To induce myofibroblasts, PDGFA containing media was replaced with media containing 10 ng/ml TGFβ (BioLegend, 763102), unless otherwise stated. For the AT1-derived ligand screen, fibroblasts in serum-free media were treated with 10ng/ml recombinant FGF1 (R&D Systems, 4686-FA), 5ng/ml WNT3A (Thermo, 315-200), 50ng/ml BMP4 (BioLegend, 595302), 50ng/ml BDNF (Thermo, 450-02) for 3 days before harvesting. For experiments involving FGF1 stimulation, FGF1 was supplemented with 10µg/ml heparin (MilliporeSigma, H4784) to stabilize the ligand and to facilitate receptor.

### Mouse AT2 isolation, culture, and differentiation

Lung dissociation was performed as described previously ^53,54^. Briefly, lungs were inflated with an enzymatic dissociation solution: 450 U/mL Collagenase I (Worthington, LS004197), 5 U/mL Dispase (Corning, 354235), and 0.33U/mL DNase I (Roche, 10104159001). Lung lobes were minced and then incubated in the enzyme solution at 37°C for 25–35 minutes, then quenched with 10% FBS/DMEM and strained. Cell pellet was resuspended in red blood cell lysis buffer (100 μM EDTA, 10 mM KHCO_3_, 155 mM NH_4_Cl) for 2 min, followed by quenching with 10% FBS/DMEM and filtration. For FACS, the cell pellet was resuspended in a sorting buffer (0.5% BSA [Genclone, 25-529F], 2 mM EDTA). Cells were stained with EpCAM/CD326-Brilliant-Violet-711 (BioLegend, 118233, 1:200), Lysotracker-Green DND-26 (Invitrogen, L7526, 1:10,000), CD140a-PE (BioLegend, 135905, 1:200), CD31-eFluor-450 (Invitrogen, 48-0311-82, 1:200), and CD45-eFluor-450 (Invitrogen, 48-0451-82, 1:200). EpCAM^+^Lysotracker^hi^ cells were collected in 2% FBS/DMEM/F12. Sorting was performed using a SONY SH800S or MA900. For differentiation, AT2s were dissociated and plated in 5% Matrigel and cultured in SFFF media for 3–5 days, then followed by 7–8 days of ADM media.

### Proximity ligation in situ hybridization

Proximity ligation in situ hybridization (PLISH) was performed as described previously ^55^. Briefly, fixed and OCT embedded mouse lungs were cryosectioned, fixed with 4% paraformaldehyde for 20 min, treated with 20 μg/ml proteinase K for 9 min at 37°C, and dehydrated with up-series of ethanol. The sections were incubated with gene-specific oligos in hybridization buffer (1M sodium trichloroacetate, 50 mM Tris (pH 7.4), 5 mM EDTA, 0.2 mg/ml heparin) for 2h at 37°C. Common bridge and circle probes were added to the section and incubated for 1h, followed by the T4 DNA ligase reaction for 2 h. Rolling circle amplification was performed by using phi29 polymerase (30221, Lucigen) for 12h at 37°C. A fluorophore-conjugated detection probe was added and incubated for 30 min at 37°C followed by mounting in DAPI-containing medium. AT2s were differentiated to AT1s as described previously ^56^ and PLISH was performed as described above.

### Immunofluorescence staining of tissue

OCT tissue sections were brought to room temperature and washed in PBS. Antigen retrieval was performed with citrate buffer, pH 6.0 (Sigma-Aldrich) using a water bath (95°C for 10–12 min). Sections were washed with 0.1% Triton in PBS (PBST), incubated in blocking buffer (1% BSA in 0.1% PBST) for 1h, and then stained with primary antibody for 2h at RT or overnight at 4°C. Following primary antibody incubation, the tissues were washed three times (5 minutes each) in PBST followed by incubation with secondary antibody in blocking buffer for 1h. Sections were washed three times with PBST (5 minutes each) and coverslips were mounted with Fluor G reagent with DAPI (Thermo Fisher Scientific, 00-4959).

### Immunofluorescence staining of cultured cells

Cultured cells were fixed in wells with 4% PFA at RT for 15 min. They were washed 3 times for 5 minutes each with PBS, permeabilized in 0.2% Triton X-100 in PBS, and incubated with 1% BSA in PBS for 30 min at RT, followed by primary antibodies at 4°C overnight. Samples were then washed 3 times in PBST, incubated with secondary antibodies for 1h at RT, washed with PBST 3 times, and incubated in 100ng/ml DAPI for 15 min (Sigma-Aldrich, D9542).

### Human lung dissociation

Human lung dissociation was performed as described previously ^54^. Briefly, 2 to 3 g of human lung tissue was washed with 1% Antibiotic-Antimycotic PBS, followed by removal of pleura, small airway and visible vasculature. The remaining tissues were minced and incubated in an enzyme mixture (Collagenase type I: 1.68 mg/ml, Dispase: 5 U/ml, DNase: 10 U/ml) at 37°C for 1-1.5h with rotation. The cells were then filtered through a 100μm strainer and rinsed with DMEM containing 10% FBS. Cell suspension was spun down at 450g for 10 min, and the pellet was resuspended in red blood cell lysis buffer for 5 min. After RBC lysis, the cells were washed with DMEM containing 10% FBS, and filtered through a 40μm strainer, then centrifuged at 450g for 5 min at 4°C. The cell pellet was then used for isolation of fibroblasts.

### Primary human fibroblasts enrichment by MACS

Fibroblast enrichment was done using magnetic activated cell sorting (MACS) as previously described ^52^. Briefly, after lung dissociation, cells were incubated in MACS buffer (1% BSA, 2 mM EDTA, antibiotic/antimycotic in PBS, pH 7.2) containing CD146 (Miltenyi Biotec 130-093-596, 1:50), CD45 (Miltenyi Biotec 130-045-801, 1:50), and CD326 (Miltenyi Biotec 130-061-101, 1:50) microbeads at 4°C for 1h with rotation. Cells were then washed and incubated with CD31 biotinylated antibody (Miltenyi Biotec, 130-110-805, 1:50) at 4°C for 10 min followed by washes and incubation with streptavidin microbeads (Miltenyi Biotec, 130-048-102, 1:10) at 4°C for 15 min. Magnetic separation of antibody bound cells was performed with Miltenyi Biotec LS columns (130-042-401). Fibroblasts collected from the flowthrough were resuspended in fibroblast growth media (advanced DMEM/F12 (Gibco, 12634-010), Antibiotic/Antimycotic (Gibco, 15240062), HEPES (Thermo Fisher Scientific, 15630080), GlutaMAX (Thermo Fisher Scientific, 35050061), B27 supplement (Thermo Fisher Scientific, 17504044), N2 supplement (Thermo Fisher Scientific, 17502048), N-acetylcysteine (Sigma-Aldrich, A9165), and 10 ng/ml PDGFA (Biolegend, 773708), plated cultured on 5% Matrigel (Corning, 354230) coated dish. To induce myofibrogenesis, PDGFA containing media was replaced with media containing 10 ng/ml TGFβ (BioLegend, 781802), unless otherwise stated. Media was changed every 2 to 3 days.

### Adenoviral infection

Adenovirus encoding Cre recombinase co-expressing GFP (Ad-Cre-GFP, SL100706) and GFP alone (Ad-GFP, SL100708) were purchased from SignaGen Laboratories and added to cultures at a 1:8000 dilution. Fibroblasts were infected by adding virus stocks directly to the culture medium. After an overnight incubation at 37°C, the infection medium was replaced with fresh culture medium.

### Inhibitor experiments

Primary mouse alveolar fibroblasts were treated with small molecule inhibitors targeting PTPN11 (also known as SHP2, a tyrosine phosphatase) (SHP099), PDGFRA (CP-673451), or EGFR (Erlotinib). All inhibitors were dissolved in dimethyl sulfoxide (DMSO) to prepare concentrated stock solutions and stored at −20°C. DMSO vehicle was added to all control conditions at equivalent volumes, with final DMSO concentrations in culture medium not exceeding 0.1%. CP-673451 (Cayman Chemical,19170) was used at 10nM or 100nM. SHP099 (MedChemExpress, HY-100388) was used at 10μM. Erlotinib hydrochloride (MilliporeSigma, SML2156) was used at 100nM, and in combination with CP-673451 (10nM) to assess potential receptor compensation. For all experiments, cells were seeded and maintained in serum-free medium (SFM) prior to inhibitor addition. CP-673451 and erlotinib were added 1h before stimulation with 10 ng/ml recombinant PDGFA; SHP099 was added 2h before stimulation. Cells were harvested at day-3 post-treatment and processed for immunofluorescence imaging as described above.

### Phospho RTK Array

To profile receptor tyrosine kinase activation in response to PDGFA stimulation, the Proteome Profiler Mouse Phospho-RTK Array Kit (R&D Systems, ARY014) was used according to the manufacturer’s protocol. Primary alveolar fibroblasts in culture were lysed in Lysis Buffer 17 (R&D Systems, 895943; 1% NP-40, 20 mM Tris-HCl pH 8.0, 137 mM NaCl, 10% glycerol, 2 mM EDTA) supplemented with protease inhibitors (10 μg/ml aprotinin, 10 μg/ml leupeptin hemisulfate, 10 μg/ml pepstatin A) and kept on ice throughout. Two hundred micrograms of total protein per condition were incubated with membranes overnight at 2–8°C following blocking. Following washing, membranes were incubated with an HRP-conjugated pan anti-phosphotyrosine antibody (1:5000), followed by chemiluminescent detection. Spot pixel density was quantified by densitometry, and signal intensity was calculated by subtracting local background from each spot. Data are presented as background-subtracted pixel density values averaged across duplicate spots.

### Image acquisition and processing

Images were captured on an Olympus FV3000 confocal microscope using 10X, 20X, 40X, and 30X objectives. Images from lung tissue staining are presented as a z-projection over approximately 3-4 z-stacks unless otherwise stated.

### Quantification of Immunofluorescence in Primary Cell Culture

#### DAPI, Ki67, and ACTA2 quantification (CellProfiler)

Immunofluorescence images from fibroblast cultures were analyzed using CellProfiler (version 5, build 427; Broad Institute) ^57^ and QuPath (version 0.6.0) ^58^. For CellProfiler analysis single-channel images were imported into a batch pipeline. Nuclei were identified from the DAPI channel using the *IdentifyPrimaryObjects* module with Global Minimum Cross-Entropy thresholding and watershed-based segmentation; objects outside the range of 20–140 pixels in diameter were excluded. Ki67 positivity was determined per nucleus using the *MeasureObjectIntensity* module applied to the Ki67 channel, with an intensity threshold applied via *FilterObjects* to classify each nucleus as Ki67-positive or Ki67-negative; results were expressed as the number and percentage of Ki67^+^ cells relative to total DAPI-counted nuclei. ACTA2 signal was quantified as total positively stained area per field using intensity thresholding and expressed as µm² of ACTA2^+^ area, normalized to total cell number were indicated throughout. Per-image measurements were exported to CSV using the *ExportToSpreadsheet* module. QuPath analyses were performed with custom batch Groovy scripts. Nuclei were detected in the DAPI channel (CH1) using the StarDist2D extension (dsb2018_heavy_augment model; detection probability threshold 0.3; nuclear diameter 40–800µm²). Ki67 (CH4) positivity was scored per nucleus using a mean intensity threshold. ACTA2 (CH2) was quantified as thresholded positive pixel area (µm²) with a minimum size filter All measurements were exported to CSV for downstream statistical analysis.

### Quantification of Immunofluorescence in Lung Tissue Sections

Lung tissue sections stained for DAPI, ACTA2, LEL, and PDGFRA/tdTomato were imaged at 20× and analyzed using QuPath (version 0.6.0) with custom batch Groovy scripts. Tissue coverage and marker-specific metrics were quantified per image as follows.

Total tissue coverage: A merged signal from all fluorescence channels was used to generate a tissue mask distinguishing lung parenchyma from background. Total tissue area (µm²) and fractional image coverage were calculated per image.

LEL^+^ coverage: The LEL channel was segmented by intensity thresholding to identify LEL-positive area.

ACTA2^+^ area and signal intensity: The ACTA2 channel was segmented by intensity thresholding to identify ACTA2-positive regions. Total ACTA2^+^ area (µm²) and mean ACTA2 fluorescence intensity within positive regions were quantified per image. Results were normalized to total tissue area where indicated.

All threshold parameters were optimized on representative images and held constant across all images within an experiment. Measurements were exported per image to CSV files for downstream statistical analysis in GraphPad Prism.

### Quantification of mean linear intercept

Mean linear intercept (MLI) analysis was performed as previously described ^21^, adapted with automated measurement in QuPath (version 0.6.0) using a custom Groovy script. Single-channel LEL-stained images of distal lung parenchyma were acquired at 20X magnification (0.3107 μm/pixel). For each image, the LEL channel was thresholded to generate a binary tissue mask, followed by morphological closing (3 pixels) to bridge gaps in septal signal, and small, enclosed regions (≤15 μm²) were filled as tissue. A horizontal grid of four evenly spaced test lines was overlaid across the full width of each image. Interalveolar chord lengths were measured directly along each grid line, and chords intersecting the image border were excluded as incomplete. Two images were quantified per mouse and averaged to yield one MLI value per mouse, with n = 3 mice per condition, and significance set at p ≤ 0.05.

### Quantification of ablation mouse models

Macrophage number was quantified using QuPath (version 0.6.0). Briefly, nuclei were detected in the DAPI channel using the StarDist2D deep learning-based nucleus segmentation model (dsb2018_heavy_augment), with a detection probability threshold of 0.15 and cell expansion of 10 µm. Detected cells were filtered by nuclear area (20–500 µm²), and F4/80 signal intensity was measured as the mean nuclear intensity per cell. Threshold was defined as the mean F4/80 intensity plus 2 standard deviations across all detected cells within that image, and cells exceeding this threshold were classified as F4/80-positive macrophages. Batch processing and CSV export were performed using a custom Groovy script in QuPath. SFTPC^+^ AT2s and TCF21^+^ alveolar fibroblasts were quantified by manual counting. For both, the total number of positive cells per field of view was recorded in ImageJ.

### Samples preparation for bulk RNA-seq and ATAC-seq

Cells were removed from culture plates with Trypsin, spun down with 10% FBS and then resuspended in PBS. ATAC-seq and RNA-seq libraries were prepared simultaneously from the same cells using the low-input ATAC & mRNA-seq protocol ^59^. Briefly, whole-cell ATAC tagmentation was performed with the Illumina Tagment DNA Enzyme and Buffer kit (Illumina, 20034198). mRNA was captured directly from tagmented cells using the Dynabeads mRNA DIRECT Micro Purification Kit (Invitrogen, 61021), followed by solid-phase first-strand cDNA synthesis with the SuperScript IV First-Strand Synthesis System (Invitrogen, 18091050). The resulting mRNA/cDNA hybrids were directly tagmented using the Nextera XT DNA Library Preparation Kit (Illumina, FC-131-1024). ATAC DNA was purified with the MinElute PCR Purification Kit (Qiagen, 28006) and amplified using the Q5® High-Fidelity 2X Master Mix (NEB, M0492). Final libraries were cleaned with DINOMAG™ Clean-Up & Size Selection SPRI Magnetic Beads (Redoxica, DN9004), quantified with a Qubit dsDNA HS Assay Kit (Invitrogen, Q32851), and assessed on an Agilent Fragment Analyzer System before paired-end sequencing on an Illumina platform.

### Bulk RNA-seq analysis

Raw paired-end reads were trimmed using TrimGalore with NextSeq quality trimming (--nextseq 20) and processed using the nf-core/rnaseq pipeline (v3.22.2) ^60^, with STAR alignment ^61^ and Salmon quantification ^62^, and aligned to the mouse reference genome mm39 using the NCBI RefSeq annotation. Differential expression analysis was performed in R using DESeq2 ^63^, with all conditions normalized simultaneously. Pairwise contrasts were computed between each treatment condition and the serum-free medium (SFM) control, as well as between Combo and each single treatment. Genes with adjusted p-value < 0.05 and |log2 fold change| > 1 were considered differentially expressed. Volcano plots were generated using EnhancedVolcano. Heatmaps were generated using pheatmap. Principal component analysis was performed using the prcomp() function in R and visualized using ggplot2. Cell-cell interaction analysis was performed using CellChat v2. ^26^

### Bulk ATAC-seq analysis

Triplicate ATAC samples from each condition were processed using the nf-core ATAC seq v2.1.2 pipeline in Nextflow v25.10.2 using a Singularity profile. Libraries were trimmed using TrimGalore 0.6.7 with quality and Nextseq trimming at q>=20. Trimmed libraries were then aligned with the mm39 genome using BWA-MEM v0.7.17 ^64^ and deduplicated using Picard v3.0.0. Deduplicated libraries were then processed and sorted using Bamtools v2.5.2 ^65^. Processed bam files were used to call peaks using MACS2 v2.2.7.1 ^66^ with narrow peak calling, FDR<0.1. Peaks were annotated to their respective genes using HOMER v4.11 ^67^. Transcription Factor motif activity was estimated from bulk ATAC sequencing using chromVar v1.32.0 ^68^ in R v4.5.2. Reference matrices were generated from the JASPAR 2020 CORE collection with human motifs, and motif occurrence in ATAC peaks was determined using “matchMotifs”. For differential motif activity, background peak sets were generated using “getBackgroundPeaks”, and motif activity deviation z scores relative to the background peak set were calculated using “deviationScores”. Pairwise comparisons for each condition against SFM were done using “differentialDeviations (paramteric = TRUE).” Differential motifs were called with p<0.05.

### CellChat analysis of ligand-receptor signaling in single cell transcriptomic data

For cell-cell communication inference, we used Seurat v5.4.0 ^69^ and CellChat v2.2.0.9001 ^70^ in R v4.5.2. Briefly, we utilized previously published single-cell datasets for control mouse mesenchyme (GSE218666, GSE173878) ^21^, control mouse epithelium (GSE158192) ^71^, normal human mesenchyme (GSE135893, GSE132771, GSE196638) ^72–74^ and normal human epithelium (GSE135893). For both mouse and human Seurat objects, we identified and removed doublets using ^72–74^ and normal human epithelium (GSE135893). For both mouse and human Seurat objects, we identified and removed doublets using ^71^, normal human mesenchyme (GSE135893, GSE132771, GSE196638) ^72–74^ and normal human epithelium (GSE135893). For both mouse and human Seurat objects, we identified and removed doublets using ^21^, control mouse epithelium (GSE158192) ^71^, normal human mesenchyme (GSE135893, GSE132771, GSE196638) ^72–74^ and healthy human epithelium (GSE135893). For both mouse and human Seurat objects, we identified and removed doublets using ^71^, normal human mesenchyme (GSE135893, GSE132771, GSE196638) ^72–74^ and normal human epithelium (GSE135893). For both mouse and human Seurat objects, we identified and removed doublets using scDblFinder v1.24.0 ^75^ and removed additional cells expressing over 15% mitochondrial content, below 500 features, or over 6,000 features. Individual datasets were then normalized using sctransform v0.4.3 ^76^, followed by PCA using “RunPCA.” Normalized datasets were prepped for integration using “PrepSCTIntegration,” anchors were found using “FindIntegrationAnchors,” and the objects were integrated using the identified anchorset. We then reran PCA and generated UMAP visualization for each integrated dataset using “RunUMAP.” For CellChat, we normalized RNA counts from the integrated objects using “NormalizeData.” We then created a cellchat object using normalized RNA counts, and kept respective cluster annotations for each population using “createCellChat.” We then limited the analysis to secreted signaling, by subsetting the expression data to signaling genes. Highly expressed ligands and ligand receptors were identified using “identifyOverExpressedGenes” and “identifyOverExpressedInteractions.” Communication probabilities were computed with “computeCommunProb.” For the mouse comparison, average gene expresseion was computed with the default trimean method, whereas the human comparison was computed as a 15% truncated mean with “type=”truncatedMean”, trim=015).” Cell-cell interactions were filtered using “filterCommunication” with “min.cells = 10.” Communication probabilities were then inferred using “computeCommunProbPathway.” Aggregate cell-cell communications were calculated using “aggregateNet.” Outgoing and incoming signal communication heatmaps were generated using “netAnalysis_signalingRole_heatmap” with pattern = “outgoing” or pattern = “incoming”. Circle plots depicting PDGFa to PDGFRa signaling were generated by using “netVisual_aggregate,” with signaling = “PDGF”. To determine the differing roles of each cell population in PDGF signaling, we used “netAnalysis_computeCentrality” with inferred intercellular communication (“netP”), and visualized using “netAnalysis_signalingRole_network.”

### Phosphoproteomics Cell Treatment and Sample Preparation

Primary mouse alveolar fibroblasts were plated on 5% matrigel-coated surfaces and grown to approximately 80% confluency. Prior to stimulation, cells were washed with PBS and serum-starved overnight in serum-free medium (SFM). Cells were given ligand stimulation as follows: SFM control (vehicle only); 80 ng/ml PDGFA for 30 min; 20 ng/ml TGFβ for 30 minutes; or combined stimulation with 20 ng/ml TGFβ for 30 min followed by 80 ng/mL PDGFA for the final 7 min. Four biological replicates were collected per condition (n = 4).

Following ligand stimulation, cells were washed twice with cold PBS. Cells were lysed on ice with RIPA buffer with Halt Protease and Phosphatase Inhibitor Cocktail (Thermo Scientific, #1861280). Lysates were sonicated briefly on ice, and protein concentration was determined by Bradford assay. Lysates were then submitted to the Duke Proteomics and Metabolomics Core Facility (DPMCF).

### Phosphopeptide Enrichment and LC-MS/MS Acquisition

Sample processing was performed by the DPMCF. Protein lysates were adjusted 5% SDS in 50 mM TEAB buffer (pH 8.5), reduced with 100 mM DTT at 80°C for 10 min, and alkylated with 250 mM iodoacetamide in the dark for 30 min. Samples were processed using S-trap micro devices (Protifi) and digested with Sequenz modified bovine trypsin (Worthington) and LysC (Worthington) at 47°C for 2h. Peptides were eluted and lyophilized prior to phosphopeptide enrichment. Phosphopeptide enrichment was performed using the KingFisher Flex protocol with MagReSyn Zr-IMAC magnetic nanoparticles in the presence of 0.25M glycolic acid. Enrichments were quenched with 10% formic acid. A study pool quality control (SPQC) sample was prepared by pooling equal volumes of all post-enrichment samples; four SPQC replicates were included and analyzed in block-randomized order alongside experimental samples.

Quantitative LC-MS/MS was performed using an Evosep One LC system coupled to a ThermoFisher Orbitrap Astral mass spectrometer via a Nanospray Flex ionization source. Peptides were separated using a 30-samples-per-day method on a Pepsep 15 cm × 150 µm column (1.5 µm particle size). MS1 precursor scans were acquired at 240,000 resolution over a mass range of 380–1080 m/z (AGC target 500%, maximum injection time 50ms, collected every 0.6s in centroid mode). Data-independent acquisition (DIA) MS/MS was performed with 5 m/z isolation windows over the same precursor range (AGC target 500%, maximum injection time 8ms, NCE 28, default charge state 3).

Raw MS data were converted to .htrms format and processed in Spectronaut 20 (Biognosys). A spectral library was generated using direct-DIA searches against the *Mus musculus* UniProt database (downloaded April 8, 2024; 17,243 entries including appended contaminant sequences). Search parameters specified trypsin/P cleavage with up to 2 missed cleavages, peptide lengths of 7–52 amino acids, variable acetylation (protein N-terminus) and phosphorylation (Ser/Thr/Tyr) modifications, and fixed carbamidomethylation (Cys). Data were filtered at a 1% precursor and protein group false discovery rate (FDR). PTM site analysis was performed with a localization probability threshold of 0.75 to retain site-localized phosphopeptides. Local normalization was applied at the phosphopeptide level, and quantitative data were aggregated to the PTM site level using linear mode site collapse.

Non-localized phosphosite data were used for downstream quantification to minimize missingness, retaining phospho PTM sites localized in at least 3 runs; redundant rows due to localization ambiguity were concatenated. This yielded 28,804 quantified phosphosites. Missing value imputation was performed by Spectronaut based on background signal. Median percent coefficient of variation for SPQC technical replicates was 13.9% in non-imputed data and 15.9% in imputed data.

### Phosphoproteomics Statistical Analysis and Visualization

Imputed PTMsite-level abundance data were log2-transformed prior to statistical testing. Paired t-tests were performed on log2-transformed data for six pairwise comparisons: PDGFA vs. SFM, TGFβ vs. SFM, combination vs. SFM, PDGFA vs. TGFβ, combination vs. PDGFA, and combination vs. TGFβ. P-values were adjusted for multiple testing using the Benjamini-Hochberg procedure. Phosphosites were considered significant at p < 0.05 and |log2 fold change| > 1. Results for all comparisons were exported and visualized using volcano plots and heatmaps. Pathway enrichment analysis was performed on significantly regulated phosphosites for each comparison. All visualization was performed using ggplot2 and pheatmap in R. We utilized the SIGNOR v4.0 ^77^ database with mouse (taxon 10090) and human (taxon 9606) to annotate phosphosites as inhibitory or activating. Phosphosites were initially aligned with the SIGNOR mouse dataset, and then remaining phosphosites were aligned with the human dataset to annotate sites curated only in the human database. Using this method, we annotated 1,044 phosphosites within our dataset.

### Kinase activity inference using benchmarKIN

Kinase activity was inferred with benchmarKIN v1.0.0 package ^78^ using the “run_zscore” function with “minsize=5”. Using this threshold, we identified 72 kinases with 1,721 kinase-substrate pairings, with CDK1 being the most abundant with 117 substrate sites. P values were computed.

### Transcription Factor Regulon scoring

We measured transcription factor regulon activity using decoupleR v2.2.0 in python v3.12 ^79^ with the argument ‘dc.mt.ulm(tval=True)’ with ‘tmin=5’ (univariate linear model with score = t-value). We used variance stabilized counts and scored regulon activity with the CollecTRI mouse network ^80^. Regulon subsets were curated manually based on gene function and respective upstream cofactors with SRF – MRTF-related genes (n= 21), TCF related genes (n= 8). E2F shared genes were identified by overlapping genes within each E2F family member regulon, and subsetting them into cell cycle genes (n= 24), and tumor suppressor genes (n = 15). Regulon activity values were plotted as the regulon mean for each condition relative to SFM. Likewise, we measured relative chromatin accessibility of the same genes within each regulon subset, by assigning peaks to each gene based on proximity to the promoter (+/− 5 kb from the transcription start site). For both contexts, shaded area represent +/− SEM, and asterisks mark p<0.05 using Student’s two sample t-test versus SFM.

### Pathway enrichment analysis

We utilized msigdbr and KEGG databases to undergo pathway enrichment for Hallmark and KEGG pathways respectively. We used gseapy 1.3.1 ^81^ with the ‘gp.prerank’ argument for each Deseq2 contrast with a threshold removing genes with baseMean <10, min_size = 15, and max_size = 500. ‘gp.enrich, hypergeometric’ was then used to identified significantly enriched pathways relative to SFM. Significance in pathway enrichment was defined as p <0.05 and Log2FC >= 0.568.

### Antibodies

Primary antibodies were as follows: mouse PDGFRA (R&D Systems, AF1062, goat, 1:250), human PDGFRA (R&D Systems, AF307NA, 1:500), α-Smooth Muscle Actin/ACTA2 (MilliporeSigma, C6198, Cy3-conjugated, 1:1000), α-Smooth Muscle Actin/ACTA2 (BioLegend, 614856, Alexa Fluor 647-conjugated, 1:1000), Ki67 (ThermoFisher, 14-5698-82, rat, 1:500), TCF21 (Invitrogen, PA5-119812, rabbit, 1:200), TdTomato (Origene, AB8181-200, rabbit, 1:700), pro-SPC (MilliporeSigma, AB3786, rabbit, 1:500), F4/80 (ThermoFisher, 14-4801-81, rat, 1:250), SFN (ThermoFisher, PA5-95056, rabbit, 1:250), RAGE/AGER (R&D Systems, MAB1179, mouse, 1:500), lycopersicon Esculentum Lectin conjugated to DyLight 649 (Vector Laboratories, DL-1178, 1:1000), and lycopersicon Esculentum Lectin conjugated to DyLight 488 (Vector Laboratories, DL-1174-1, 1:1000).

Secondary antibodies were all used at 1:500 dilution and were as follows: Alexa Fluor 594 donkey anti-goat IgG (ThermoFisher, A11058;), Alexa Fluor 488 donkey anti-goat IgG (ThermoFisher, A11055), Alexa Fluor 488 donkey anti-rabbit IgG (ThermoFisher, A21206;), Alexa Fluor 647 donkey anti-rabbit IgG (ThermoFisher, A31573), Alexa Fluor 647 donkey anti-goat IgG (ThermoFisher, A21447), Alexa Fluor 488 donkey anti-rat IgG (ThermoFisher, A21208), Alexa Fluor 594 donkey anti-rat IgG (ThermoFisher, A21209), Alexa Fluor 594 goat anti-rabbit IgG (ThermoFisher, A21207), Alexa Fluor 647 goat anti-rat IgG (ThermoFisher, A21247), Alexa Fluor 647 goat anti-rabbit IgG (ThermoFisher, A21245).

### Statistical analysis

Statistical methods relevant to each figure are outlined in the figure legend. Sample size was not predetermined. All experiments were performed on at least three biological replicates. Data are presented as mean ± SD unless otherwise indicated in the figure legend. Statistical analysis was performed in GraphPad Prism and R. An unpaired two-tailed Student’s t-test was used for comparisons between two conditions. Two-way ANOVA with Šídák’s multiple comparisons test was used where two factors were compared. One-way ANOVA followed by Dunnett’s multiple comparisons test was used for comparisons against a single control condition, and Tukey’s multiple comparisons test where all pairwise comparisons were made. Statistical significance was set a priori at p < 0.05. Statistical methods for bulk RNA-seq, bulk ATAC-seq, and phosphoproteomic analyses are described in their respective methods sections above.

## Supporting information

All Supplemental Figures

## Data Availability

Bulk RNA-seq and ATAC-seq data generated in this study have been deposited in the NCBI Gene Expression Omnibus (GEO) and are publicly available under accession numbers GSE348864 and GSE348865, respectively. Phosphoproteomic data has been deposited in PRIDE database under accession number PXD084815.

## Acknowledgements

We thank Preetish Kadur Lakshminarasimha Murthy and Vishwaraj Sontake for help with initial experimental set up as well as Bailey Layish and Lucas McNeill for technical support. We thank Tata lab members for fruitful discussions, and the Duke Compute Cluster for server space and data storage. We thank Dr. Michael Dee Gunn for kindly providing *CD169-DTR* mice and Dr. Johanna Andrae for kindly providing *Pdgfa^fl/fl^* mice. We thank the Duke University School of Medicine for the use of the Proteomics and Metabolomics Core Facility, which provided quantitative phosphoproteomic analysis by data-independent acquisition (DIA) mass spectrometry. This work is the result of NIH funding, in whole or in part, and is subject to the NIH Public Access Policy. Through acceptance of this federal funding, the NIH has been given a right to make the work publicly available in PubMed Central. A.K. is supported by a fellowship from NIH/NHLBI (F30HL143911). T.W. received funding support from Duke pulmonary research training grant (PROSPER) 5T32HL160494-05. This work is supported by NHLBI/NIH R01HL160939 and R01HL153375, and a pilot grant support from NCI/NIH P30 Cancer Center Support Grant (P30CA014236) to P.R.T R01HL174525 to A.T. This work was partially supported by funds from Whitehead foundation; P.R.T. is a Whitehead Scholar at Duke University.

## Author contributions

R.S.F. co-designed and performed ex vivo and in vivo experiments, analyzed data, prepared figures, and co-wrote the manuscript. A.K. co-designed and performed in vivo experiments. T.W. analyzed ATAC-seq and transcriptome profiling, assisted in computational analysis. K.T. processed samples for bulk RNA-seq and ATAC-seq analysis. M.F. assisted with Phospho-proteome profiling experimental design. S.L. performed in vivo experiments, acquired images, and analyzed data. A.T. co-designed, and performed in vivo experiments, performed image acquisition, and prepared figures. P.R.T. conceived, co-designed, supervised the work, and wrote the manuscript. All authors reviewed and edited the manuscript.

