## Supplementary material for "A tonic PDGFA-dependent niche organizer maintains alveolar homeostasis": All Supplemental Figures

### Extended Data Figure 1

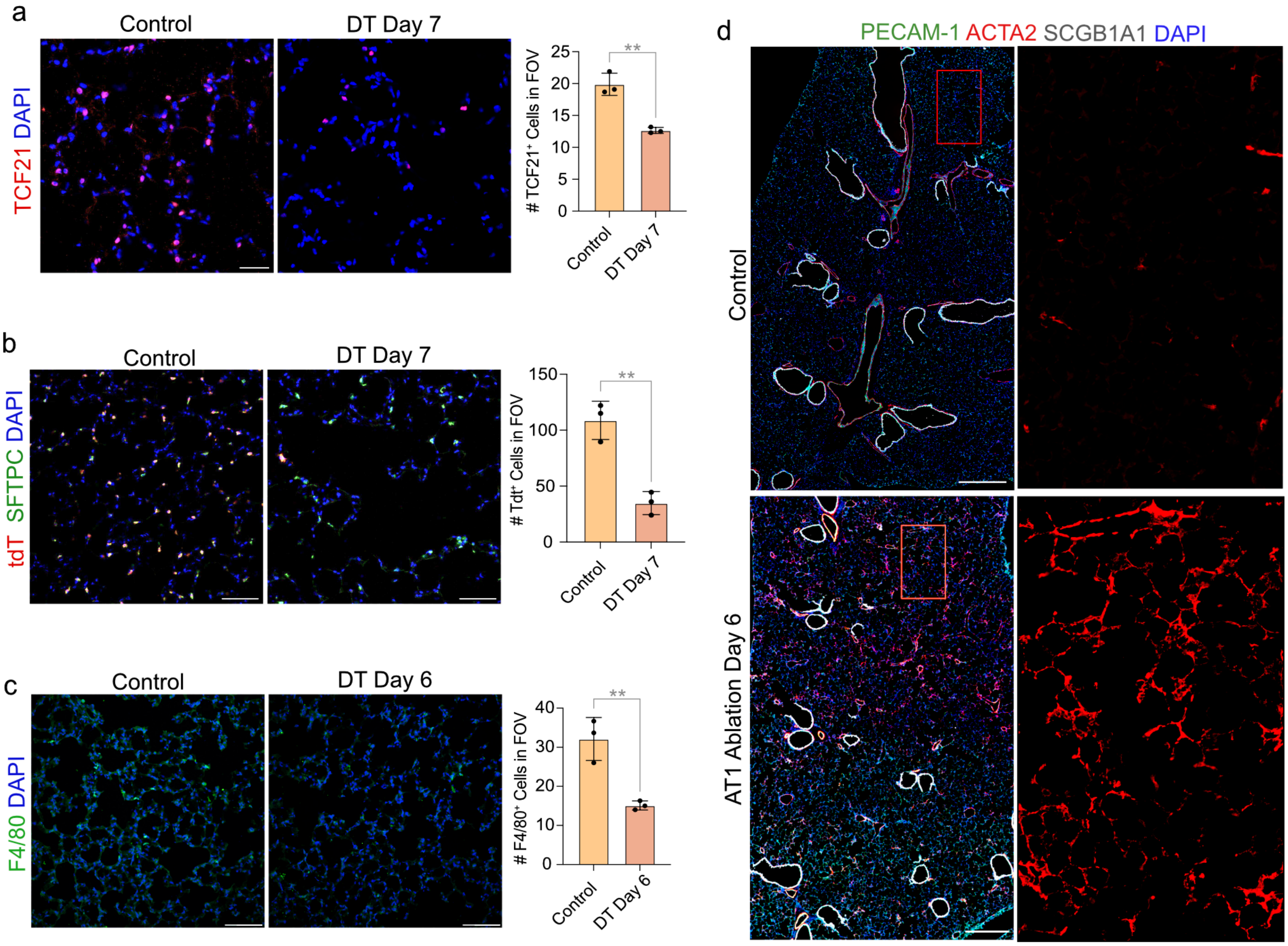

#### Extended Data Figure 2

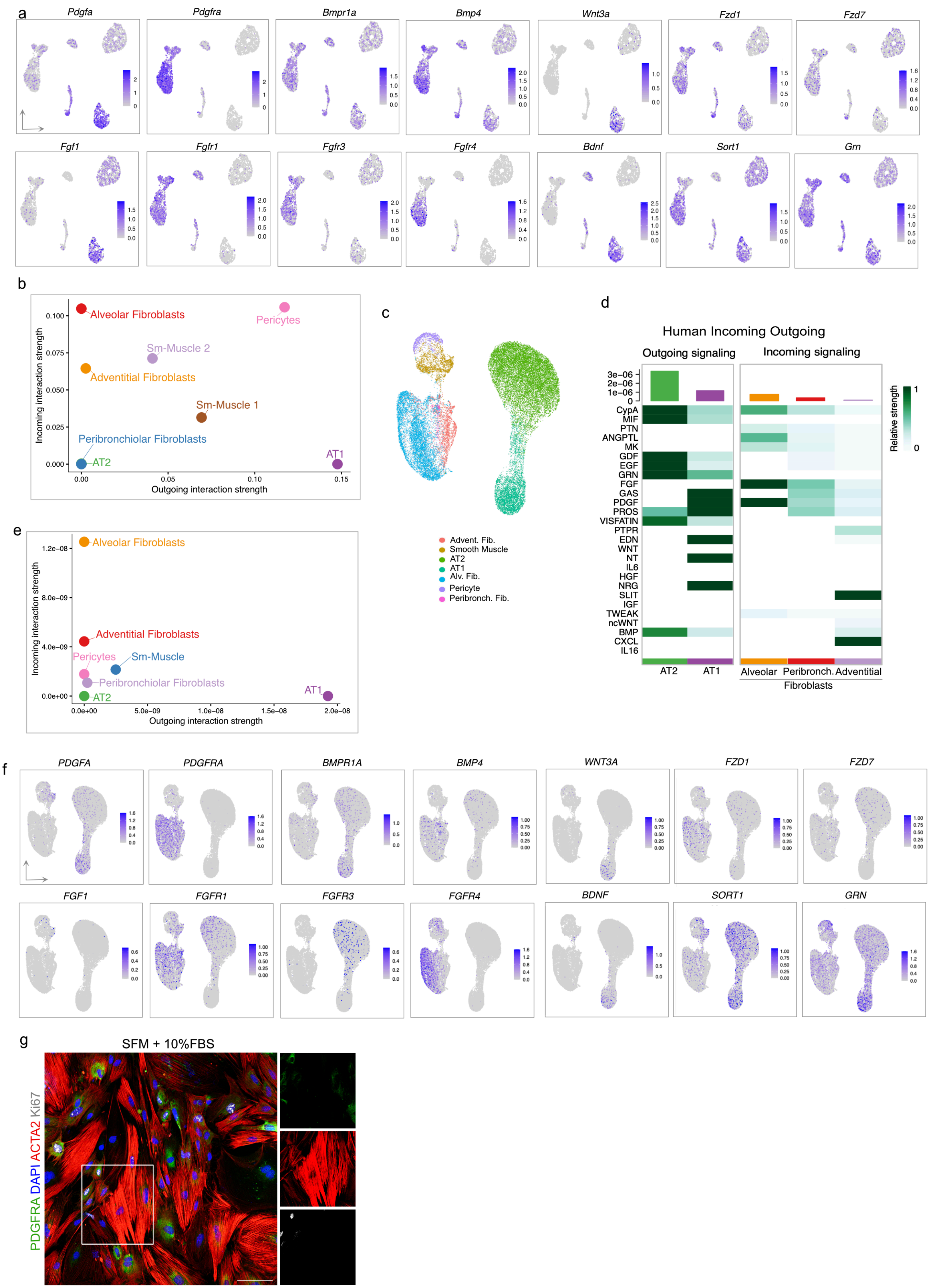

### Extended Data Figure 3

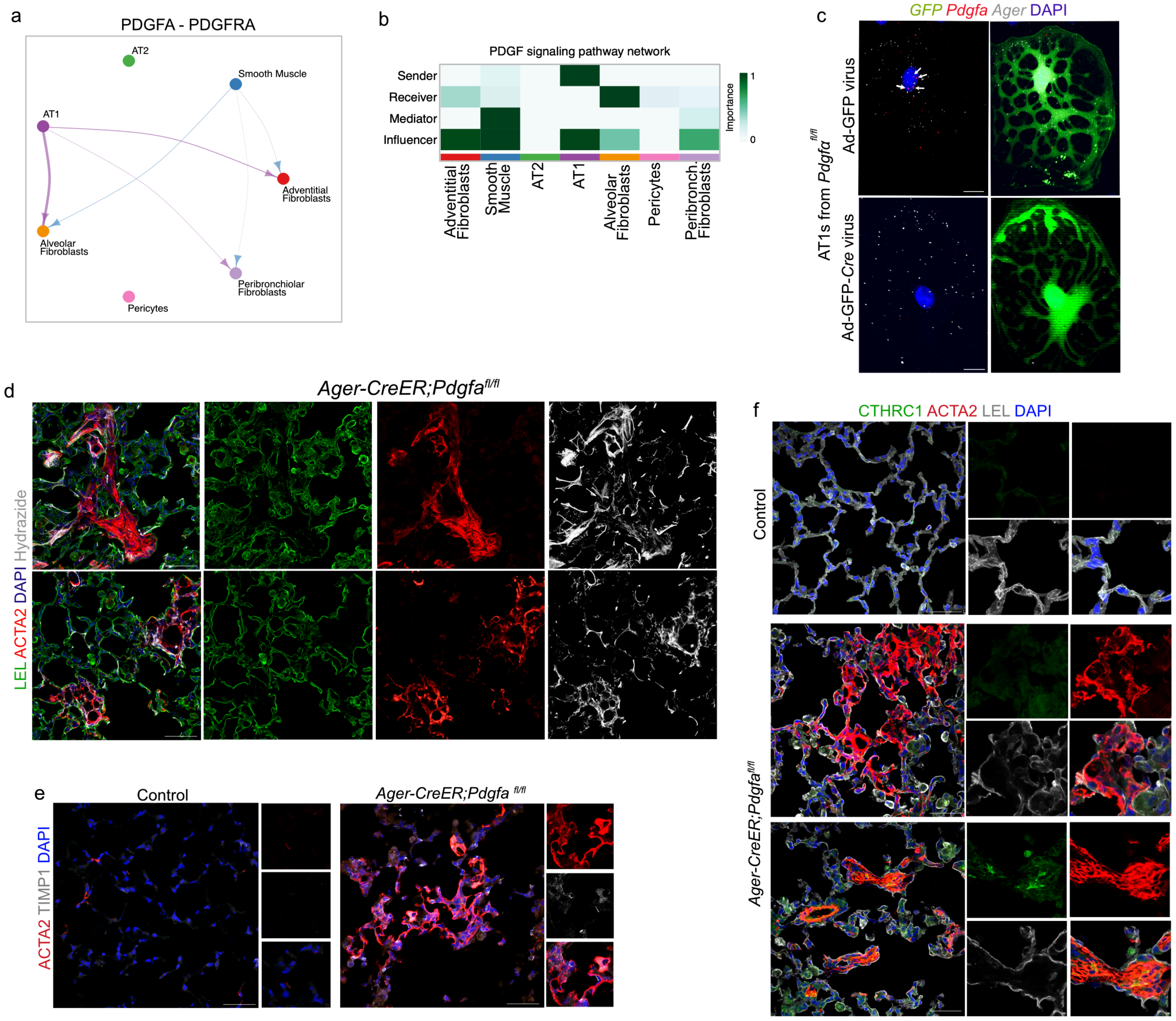

### Extended Data Figure 4

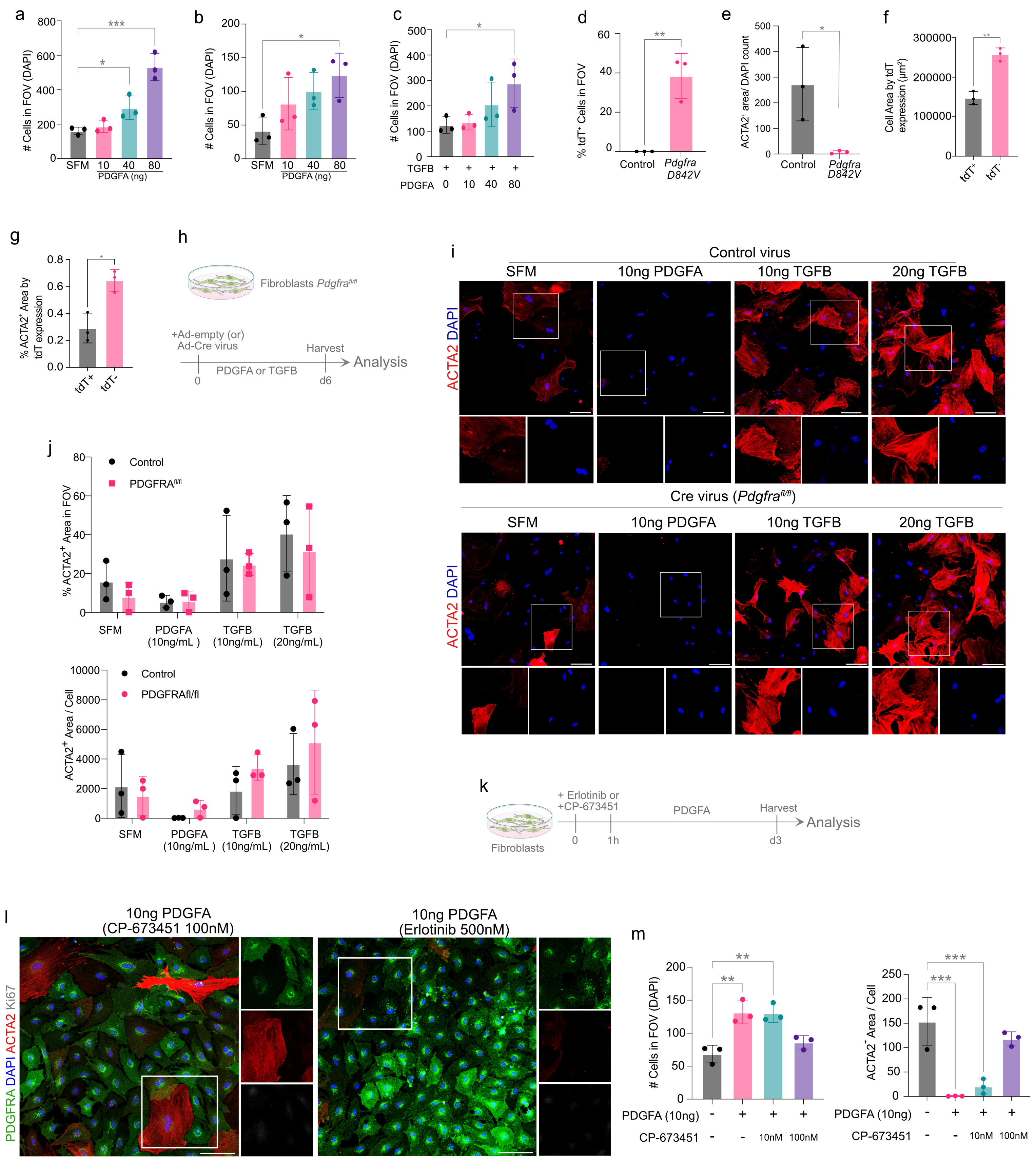

### Extended Data Figure 5

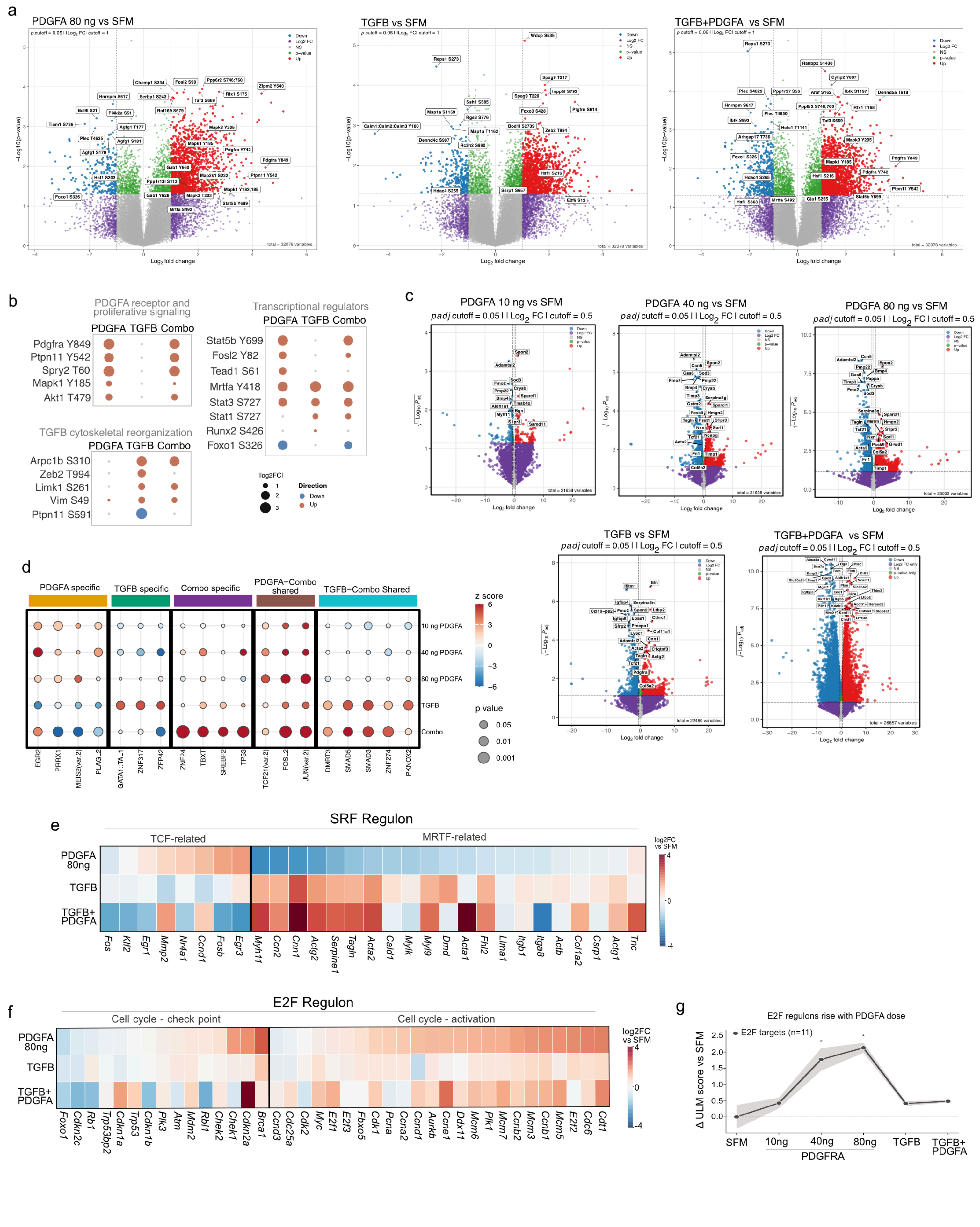

### Extended Data Figure 6

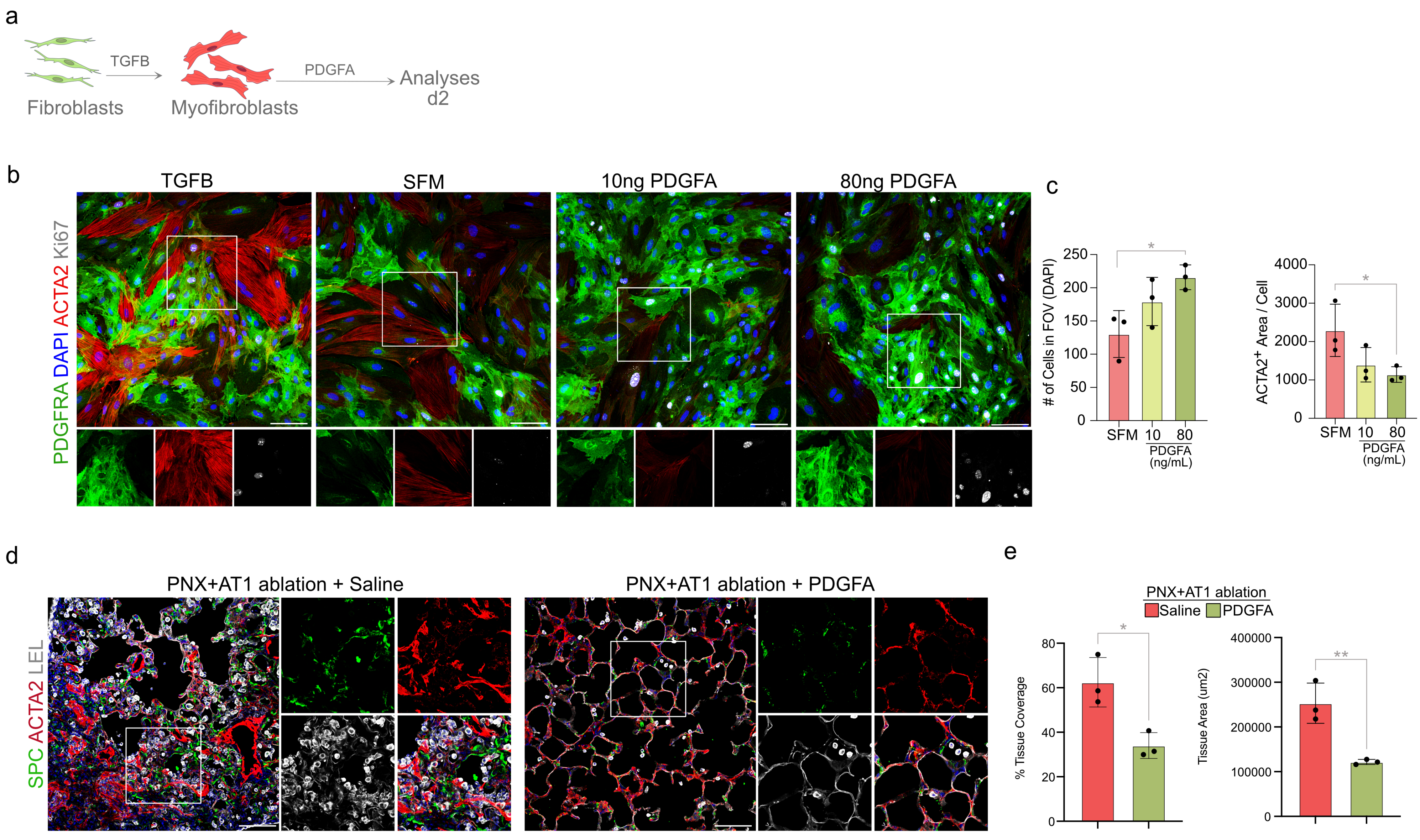
